# Parasitic connections: a patescibacterial epibiont, its methylotrophic gammaproteobacterial host and their phages

**DOI:** 10.1101/2024.03.08.584096

**Authors:** Feriel Bouderka, Purificación López-García, Philippe Deschamps, Yifan Zhou, Mart Krupovic, Ana Gutiérrez-Preciado, Maria Ciobanu, Paola Bertolino, Gwendoline David, David Moreira, Ludwig Jardillier

## Abstract

Patescibacteria form a very diverse and widely distributed phylum of small bacteria inferred to have an episymbiotic lifestyle. However, the prevalence of this lifestyle within the phylum and their host specificity remain poorly known due to the scarcity of cultured representatives. Here, we describe a complex system consisting of a patescibacterium, its gammaproteobacterial hosts, and their respective phages based on enrichment cultures and metagenomic data from two shallow, geographically close, freshwater ecosystems. The patescibacterium, *Strigamonas methylophilicida* sp. nov., defines a new genus within the family Absconditicoccaceae. It grows as epibiont on cells of methanotrophic species of the gammaproteobacterial family Methylophilaceae. *Strigamonas* cells grow tightly attached to the host, sometimes forming stacks that connect two host cells. Despite a surprisingly large genome (1.9 Mb) compared to many other Patescibacteria, *S. methylophilicida* lacks many essential biosynthetic pathways, including the complete biosynthesis of phospholipids, amino acids, and nucleic acids, implying a dependence on the host to obtain these molecules. We also identified and assembled the complete genomes of one patescibacterial phage that might represent a new virus family within the class *Caudoviricetes*, and two Methylophilaceae phages predicted to have head-tailed and filamentous virions, respectively. The patesciphage uses a modified genetic code similar to that of its host and encodes four tRNA genes, including the suppressor tRNA gene for the UGA stop codon, which is reassigned to glycine in many Patescibacteria. Our results confirm a prevalent episymbiotic lifestyle in Absconditicoccaceae and further suggest a clade-specific adaptation of this patescibacterial family for gammaproteobacterial hosts.

**IMPORTANCE:** Patescibacteria are ultra-small bacteria with reduced genomes that rely on symbiotic interactions with other prokaryotes, yet their host specificity and associated viral parasites remain poorly characterized due to limited cultured representatives. Here, we describe *Strigamonas methylophilicida*, a new patescibacterial species of the family Absconditicoccaceae that grows as an epibiont on various methylotrophic Gammaproteobacteria. This expands the host range for this family beyond previously documented photosynthetic partners. Using enrichment cultures and metagenomics, we retrieved complete genomes of novel phages infecting *S. methylophilicida* and its hosts, including one phage that uses a modified genetic code matching that of the patescibacterium. These findings reveal a previously unrecognized patescibacteria– methylotroph–phage tripartite interaction in freshwater environments, highlight the adaptations of patescibacterial phages, and shed light on the complex ecology and evolution of host-parasite-phage dynamics in understudied bacterial lineages.

## INTRODUCTION

The recently described Candidate Phyla Radiation (CPR), now formally classified as the phylum Patescibacteria (1), represents an extremely phylogenetically diverse group of bacteria, estimated to encompass between 15 and 50% of all bacterial diversity and to be present in all kinds of environments, especially suboxic ones (1–5). Patescibacteria have mostly been characterized using genome-resolved metagenomic approaches, which consistently found reduced genomes coding for limited metabolic capabilities (2, 6, 7). Notably, these bacteria lack the typical biosynthetic pathways for amino acids, nucleotides, and membrane phospholipids. Most likely, they obtain these essential molecules from other cells, which suggests that a symbiotic, likely parasitic, lifestyle is widespread within this phylum (2, 7, 8). Microscopy observations have shown that patescibacterial cells are very small and usually attached to larger cells, which supports an episymbiotic lifestyle (2, 6, 7). However, a recent report suggests that free-living patescibacterial cells might occur in freshwater samples (9), although whether these represent actual free-living stages in a parasitic lifecycle, actual free-living representatives, or dispersing stages remains to be established. Cultivation of patescibacterial representatives is therefore crucial to confidently determine the lifestyle of these bacteria.

Only a few cultured patescibacterial species have been characterized, which confirm the parasitic lifestyle inferred from genome-based approaches. The first patescibacterium ever cultured, *Candidatus* Nanosynbacter lyticus, belongs to the class Saccharimonadia and is an epiparasite of *Schaalia odontolytica* (formerly *Actinomyces odontolyticus*) in the human oral microbiome (10). Other subsequently cultured Saccharimonadia are also parasites of actinobacterial hosts (11–13). Recently, two other patescibacterial representatives have been described, *Vampirococcus lugosii* and *Absconditicoccus praedator*, representing two distant genera in the family Absconditicoccaceae (14, 15). They were both identified in saline lakes, as episymbionts of anoxygenic photosynthetic bacteria belonging to the Chromatiales (Gammaproteobacteria, within the Pseudomonadota). These species exhibit a predatory (parasitoid) lifestyle, leading to the rapid death of their hosts (14, 15). Therefore, all patescibacterial representatives characterized so far appear to be obligatory parasitic episymbionts strictly depending on hosts that are negatively impacted by their patescibacterial episymbionts. Although they are able to establish interactions with a broad range of monoderm and diderm bacteria, the two patescibacterial groups with cultured members appear so far to be specialized in interacting with hosts from specific lineages: Nanosynbacteraceae (class Saccharimonadia) parasitize Actinomycetota and Absconditicoccaceae parasitize photosynthetic Gammaproteobacteria. Some microscopy studies suggest that other Patescibacteria can apparently establish interactions also with eukaryotic and archaeal hosts (16–19). The limited availability of cultured representatives has also hampered the identification of patescibacterial phages. A few studies have established links with phage genomes using CRISPR spacers found in patescibacterial genomes. For example, Paez-Espino et al. identified 17 putative patescibacterial phages (20), Roux et al. identified 9 filamentous phages associated with Patescibacteria (21), and Liu et al. recently reported the identification of 391 additional phage genomes (22). Further efforts are required to assess the diversity and impact of phages on Patescibacteria. However, the completeness of these phage genomes varies, and none has been officially classified thus far.

In this study, we have characterized a new patescibacterial species, *Strigamonas methylophilicida* (family Absconditicoccaceae), from small freshwater ecosystems in southern Paris. It parasitizes members of the Methylophilaceae, and therefore represents the first reported association between a patescibacterium and a non-photosynthetic gammaproteobacterial host. We additionally identified several phages likely targeting *S. methylophilicida* and its hosts, one of them likely representing a novel virus family within the class *Caudoviricetes*. Our work highlights intricate parasitic interactions between cells and viruses in shallow freshwater ecosystems.

## RESULTS AND DISCUSSION

### Identification of an Absconditicoccaceae-Methylophilaceae consortium

A previous multiannual study of geographically close small freshwater ecosystems in a forest area near Paris showed that Patescibacteria constituted a large relative proportion of the prokaryotic community (23). In two of these ecosystems, the Ru Sainte Anne brook (RSA) and the Mare Gabard pond (MG), the methylotrophic bacterial families Methylophilaceae and Methylococcaceae (Beta– and Gammaproteobacteria, respectively, both now classified in the phylum Pseudomonadota (24)) were also relatively abundant depending on the date (Fig. S1). Therefore, they represented potential hosts for some of the co-occurring Patescibacteria. To enrich potential bacterial parasites relying on methylotrophic bacteria, we collected water samples from these ecosystems that we supplemented with 100 mM methanol. After one month of incubation, we observed motile bacterial cells carrying much smaller cells attached to them. We further enriched these consortia by inoculating tubes of modified nitrate mineral salts (NMS) medium with this initial enrichment to promote the growth of methylotrophic bacteria (25). After one week, we observed motile cells exhibiting curved-rod cell shape and containing numerous refringent inclusions (Fig. 1A-D), similar to described Methylophilaceae species (26). Many of these cells presented smaller epibiotic cells strongly attached to them, often forming short piles of two to four cells (Fig. 1A-D and Supplementary Video S1), which sometimes bridged two different host cells (Fig. 1C-D). These characteristics were similar to those previously described for Absconditicoccaceae species within the Patescibacteria (14, 15). We collected and fixed consortia to prepare thin sections that were examined using transmission electron microscopy (TEM) (Fig. 1E-I). The host cells were ∼2.5 µm long and ∼1.25 µm in diameter, whereas the epibiont cells were ∼300 nm in diameter and slightly flattened (n=20). The host and epibiont cells appeared tightly connected, with a gap of less than ∼20 nm separating them. The ribosomes were positioned at the periphery of the cells, whereas a dense central area probably corresponded to highly compacted DNA (Fig. 1I) (6, 15). The epibiont cell size, cell envelope structure, close attachment, and internal cell organization were consistent with those described for Patescibacteria such as *Absconditicoccus praedator* (15). However, in contrast with previous descriptions, we observed numerous extracellular vesicles of ∼20 nm in diameter around the epibiotic cells (Fig. 1E-I) which, to our knowledge, have not been previously reported in Patescibacteria.

**FIG 1.**
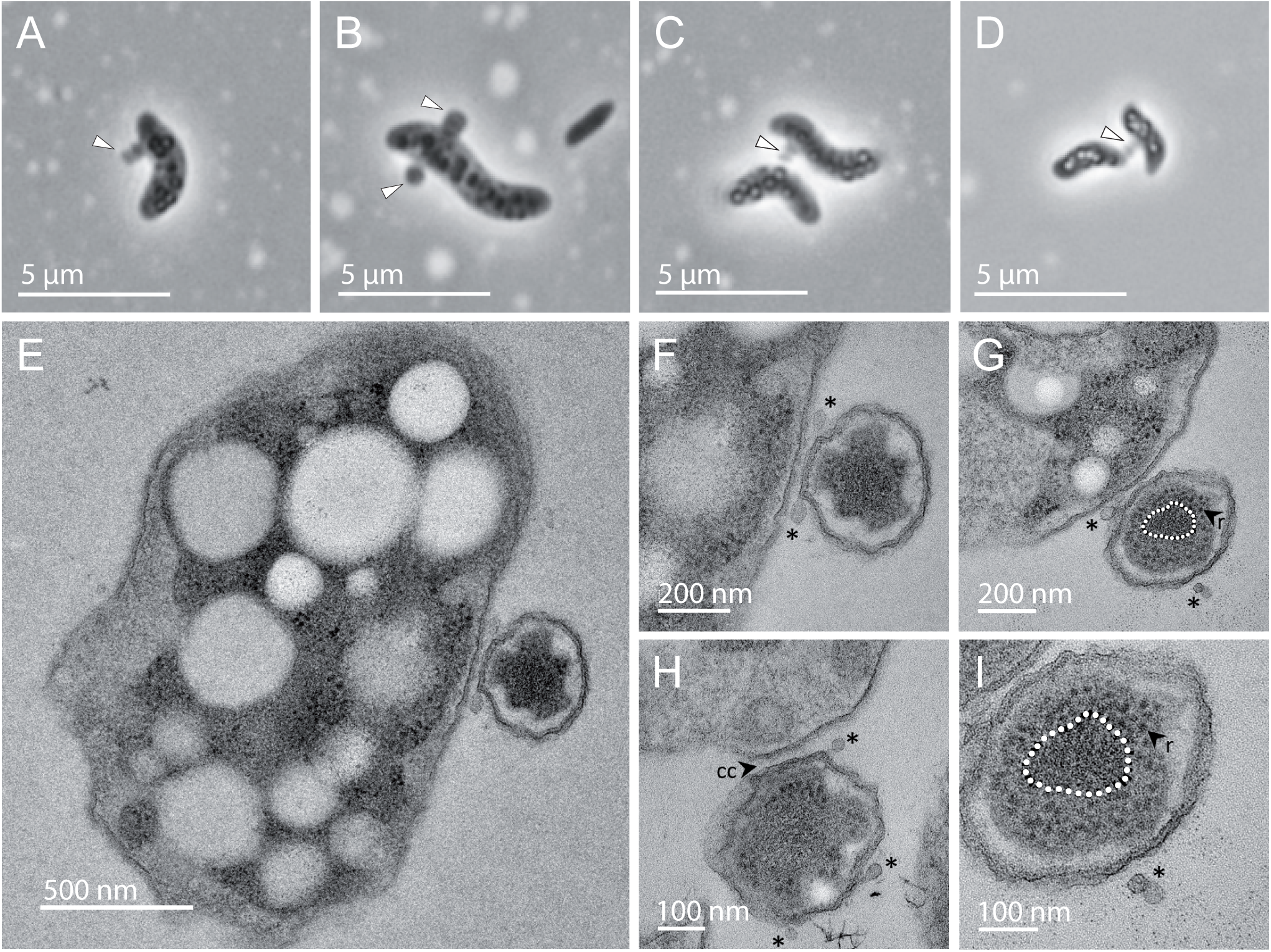
Microscopy photographs of patescibacterial-like cells and their hosts. (A-D) Phase contrast light microscopy pictures, notice the intracellular refringent inclusions in the host cells in C and D. White arrows indicate epibiotic patescibacterial cells. (E-I) Transmission electron microscopy (TEM) pictures of thin sections of the patescibacterial cells and their hosts. Asterisks indicate extracellular vesicles, CC: cell-to-cell contact, r: ribosomes. The white dashed line in panels G and I highlights the central part of the cell with dense material that probably corresponds to DNA.

Since our initial enrichments contained many other bacteria, we used fluorescence-activated cell sorting (FACS) to further enrich the identified consortia from both MG and RSA samples. FACS allowed us to sort cell populations highly enriched in these consortia (Fig. S2), as determined by microscopy observation. To identify the members of these consortia, we used a micromanipulator coupled to an inverted microscope to pick 15 individual consortia from one RSA sorted population and 15 from one MG sorted population. We then carried out single-consortium 16S rRNA gene PCR amplification. 16 consortia (10 from RSA and six from MG) yielded 16S rRNA gene amplicons, which were cloned and sequenced. In each of these consortia we identified a mix of sequences affiliated to the families Absconditicoccaceae and Methylophilaceae (Fig. S3). Regardless of the ecosystem of origin, the Absconditicoccaceae sequences were almost identical (>99%) in the 16 consortia. By contrast, we retrieved several different Methylophilaceae sequences related to the genera *Methylotenera*, *Methylovorus*, and *Methylophilus* (Fig. S3). Combined with microscopy observations, these results strongly indicated a novel type of Patescibacteria-host interaction established between Absconditicoccaceae and Methylophilaceae, where closely related Absconditicoccaceae phylotypes appear to be able to interact with hosts belonging to different Methylophilaceae genera.

To better understand this interaction using genomic data, we carried out whole genome amplification (WGA) from five FACS-sorted samples (three from RSA and two from MG) and Illumina-sequenced the amplified DNA. These mini-metagenomes contained sequences from Absconditicoccaceae and Methylophilaceae species along with some minor contaminants. We obtained two high-quality Absconditicoccaceae metagenome-assembled genomes (MAGs) by co-assembling the mini-metagenomes from RSA and MG FACS-sorted cells (see Materials and Methods). These two patescibacterial MAGs (Smet_RSA and Smet_MG, respectively) showed an average nucleotide identity (ANI) of 94.82% and an average amino acid identity (AAI) of 93.51% between them. The Smet_RSA MAG was 1.94 Mb in size with 81.1% completeness and 2.1% redundancy, while the Smet_MG MAG was 1.93 Mb with 83.1% completeness and 1.41% redundancy (Table S1). These completeness values are typically calculated via the identification of single copy genes present in most bacteria (27). Because Patescibacteria commonly exhibit genome reduction and multiple gene losses, a completeness higher than 70% is considered to correspond to complete and high quality genomes for these organisms (28, 29). The GC content of the Smet_RSA and Smet_MG MAGs was 28.47% and 27.01%, respectively. Due to the presence of multiple Methylophilaceae species in our samples, sequence binning and MAG reconstruction for the hosts were more challenging. We therefore co-assembled all samples from both ecosystems to increase genome coverage, which yielded four good-quality Methylophilaceae MAGs after two rounds of manual refinement (completeness between 90.14% and 95.77% and redundancy between 2.82% and 5.63%) (Table S1).

A maximum likelihood (ML) phylogenetic tree based on 14 concatenated ribosomal proteins revealed that the four MAGs belonged to different genera within the family Methylophilaceae (Fig. S4). For the Smet_RSA and Smet_MG epibiont MAGs, we reconstructed an ML tree using a subset of 67 markers well represented in Patescibacteria out of the 120 bacterial markers described in (1). These MAGs branched together within the family Absconditicoccaceae, forming a sister group to *V. lugosii* and*. A. praedator*, the two previously characterized representatives of this family (Fig. 2). The 16S rRNA gene sequences of our new epibionts were, respectively, 81% and 84% identical to those of *V. lugosii* and *A. praedator*. Their corresponding average amino-acid identities (AAI) were 45% and 46%, respectively. Because of low Relative Evolutionary Divergence (RED) values (0.715 for Smet_RSA and 0.716 for Smet_MG), the GTDB-tk pipeline (30) did not assign our MAGs to any known genus. Altogether, these metrics supported that our freshwater patescibacterial MAGs defined a novel genus and species (1, 31–33), which we propose to name *Strigamonas methylophilicida* (see Taxonomic appendix below).

**FIG 2.**
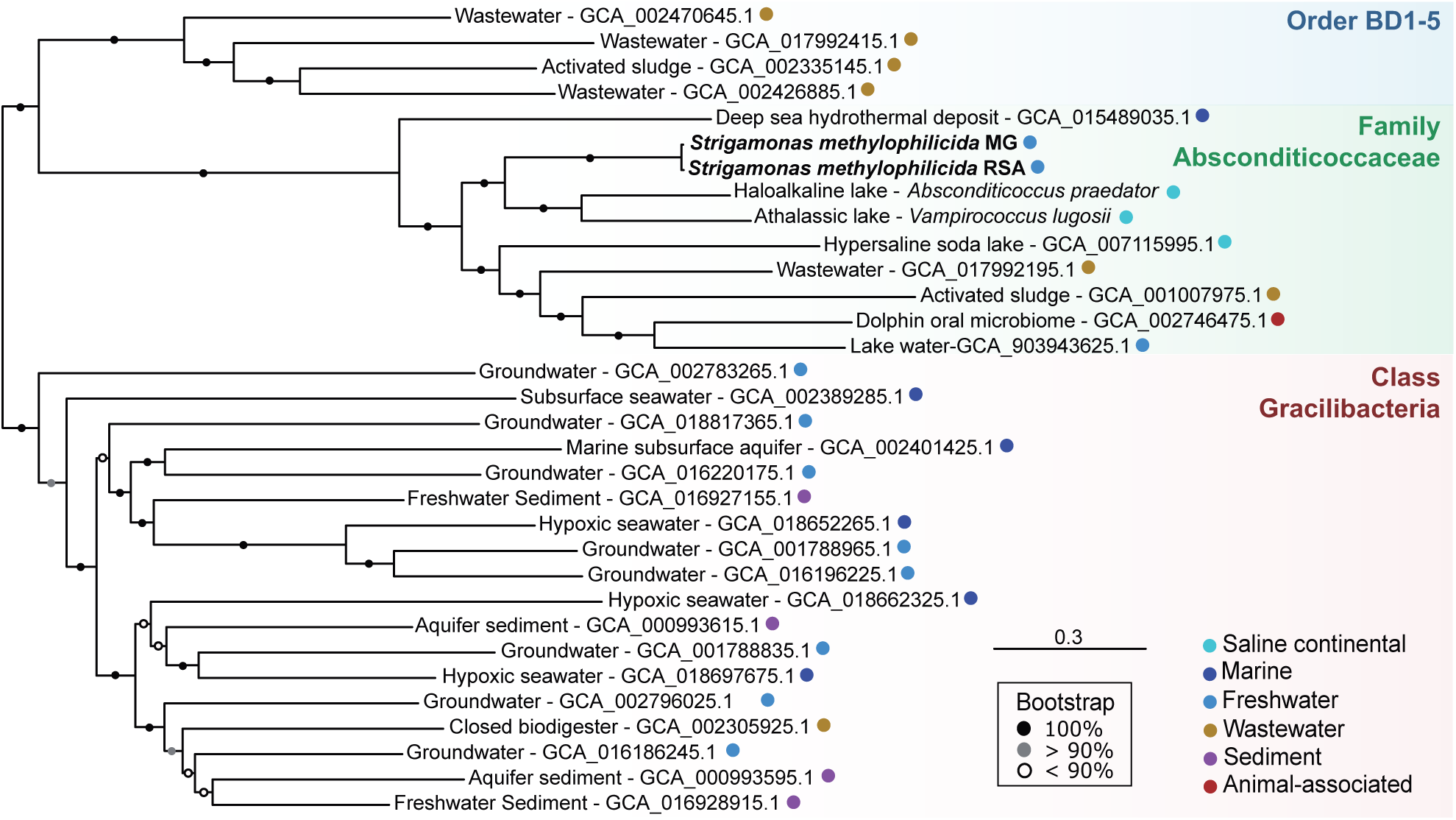
Phylogenetic placement of *Strigamonas methylophilicida* strains within the Patescibacteria. Maximum likelihood tree based on the concatenation of 67 single-copy protein markers (20,766 sites). The scale bar indicates the number of substitutions per site. Support values are based on 1000 ultrafast bootstrap replicates. Taxonomy corresponds to the Genome Taxonomy Database release 08-RS214. The environmental origin of the different taxa is indicated with colored circles next to their names.

### Gene content in Strigamonas methylophilicida

The *S. methylophilicida* Smet_RSA and Smet_MG MAGs had a size of approximately 1.9 Mb, almost twice the average genome size of most Patescibacteria (∼1 Mb, (8)). As described for other Absconditicoccaceae, *S. methylophilicida* uses a modified genetic code (code 25) where the stop codon UGA is translated to glycine (27). We identified 1,655 and 1,629 coding sequences (CDS) in the two MAGs, notably more than in other well-characterized Absconditicoccaceae representatives (1,054 in *A. praedator* and 1,124 in *V. lugosii*). However, we could only functionally annotate around one third of these CDS (565 for Smet_RSA and 543 for Smet_MG), compared to the ∼50% functionally annotated in *V. lugosii* and *A. praedator* genomes (14, 15). Although a fraction of the unannotated genes may correspond to known genes that have diverged beyond sequence similarity recognition, *S. methylophilicida* seemed to bear a substantial amount of gene novelty. The pangenome of the Absconditicoccaceae has been reported to be highly dynamic, with a very small core genome consisting of only 390 genes (14). This was even more pronounced when *S. methylophilicida* was included, revealing a mere 236 genes shared among the three genera, *Strigamonas*, *Absconditicoccus*, and *Vampirococcus* (Fig. 3a). As expected, the majority of these core genes were involved in house-keeping functions such as translation, transcription, and replication (Fig. 3b). By contrast, more than 50% of the genus-specific genes had unknown functions, and the rest were mostly involved in cell wall, membrane, and envelope biogenesis. Among the three Absconditicoccaceae genera, *Strigamonas* stood out for its substantial divergence in gene content. In addition to the 236 core genes, it shared only 34 genes with *Absconditicoccus* and 18 with *Vampirococcus*. By contrast, the two *Strigamonas* MAGs shared 1,014 genes that were not found in other Absconditicoccaceae, as well as 320 (Smet_RSA) and 268 (Smet_MG) strain-specific genes also absent in other Absconditicoccaceae (Fig. 3a). Many of these genes were organized in clusters in the two MAGs (Fig. S5) and had unknown functions (Fig. 3b, Fig. S6). However, given their very narrow distribution, at least some of them might be involved in particular ecological niche adaptations. In both MAGs, the most represented categories included defense mechanisms, in particular restriction-modification systems, protection against oxidative stress (e.g., MutT, see below), and ABC-type multidrug transport systems (Fig. 3b, Fig. S6, Table S2).

**FIG 3.**
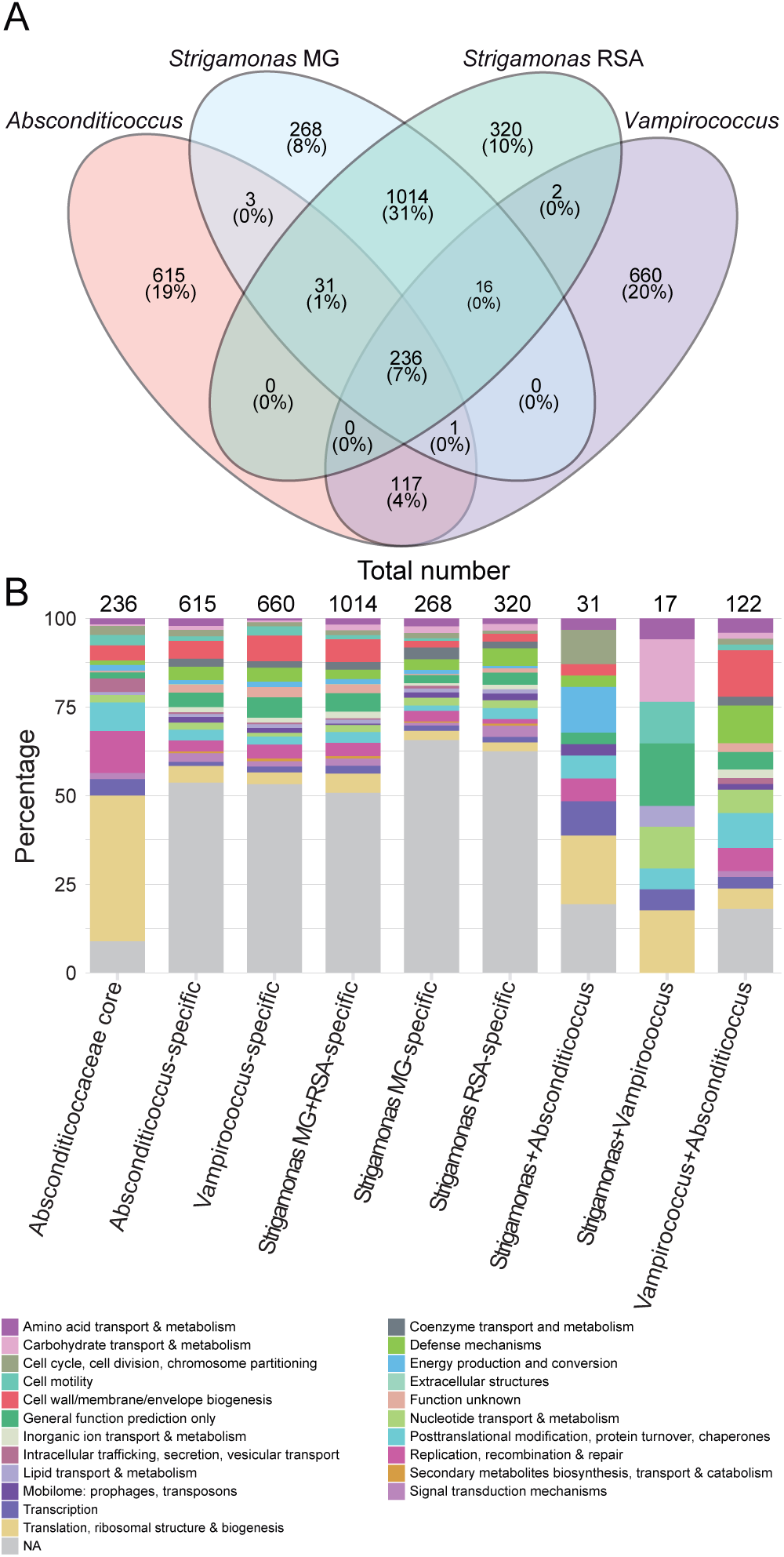
Comparison of gene content in characterized Absconditicoccaceae species. (A) Venn diagram showing the number of genes shared by different well-characterized Absconditicoccaceae species (absolute numbers and % of the total genes in the four Absconditicoccaceae species). (B) Functional annotation (COG categories) of the genes shared by these Absconditicoccaceae species. NA: not annotated.

Genes coding for most ribosomal proteins were present (n=51), with only a few missing, such as *rpL30*, which is often absent in other symbiotic bacteria (2, 8). As previously described for many other Patescibacteria, *S. methylophilicida* lacked classical phospholipid biosynthesis genes, indicating its inability to synthesize membrane lipids. Similarly, biosynthetic pathways for nucleic acids, cofactors, and many canonical amino acids were incomplete. These absences suggest strict dependence on the host for cell integrity maintenance, growth and division (7).

### Genome-based inference of the metabolism and symbiotic lifestyle

The gene repertoire of *S. methylophilicida* suggested reduced carbon and energy metabolisms. As observed previously in *V. lugosii* and many other Patescibacteria (7, 14), *S. methylophilicida* possessed an incomplete glycolysis pathway, encoding only for enzymes involved in the steps from 3-phosphoglycerate to pyruvate (Fig. 4). Therefore, these organisms most likely require 3-phosphoglycerate uptake for ATP production. *Vampirococcus* was hypothesized to obtain it from its photosynthetic host, which produces 3-phosphoglycerate via CO_2_ fixation by the enzyme ribulose 1,5-bisphosphate carboxylase/oxygenase (RuBisCO) (14). Likewise, *S. methylophilicida* might obtain 3-phosphoglycerate from its methylotrophic host (Fig. S7). Unexpectedly, akin to the photosynthetic host of *V. lugosii*, two of our Methylophilaceae MAGs encoded enzymes of the Calvin-Benson-Bassham (CBB) cycle, including phosphoribulokinase (PRK) and RuBisCO (Fig. S8). The genes encoding these two enzymes shared the same genomic context in the Methylophilaceae MAGs, being close to the CO_2_-responsive transcriptional regulator CbbR (Fig. S9). Another Methylophilaceae MAG encoding a complete CBB cycle has been recently reported (34), supporting that various Methylophilaceae can, at least facultatively, fix carbon and synthesize 3-phosphoglycerate using various pathways. Interestingly, all hosts of Absconditicoccaceae epibionts identified to date possess RuBisCO (14, 15).

**FIG 4.**
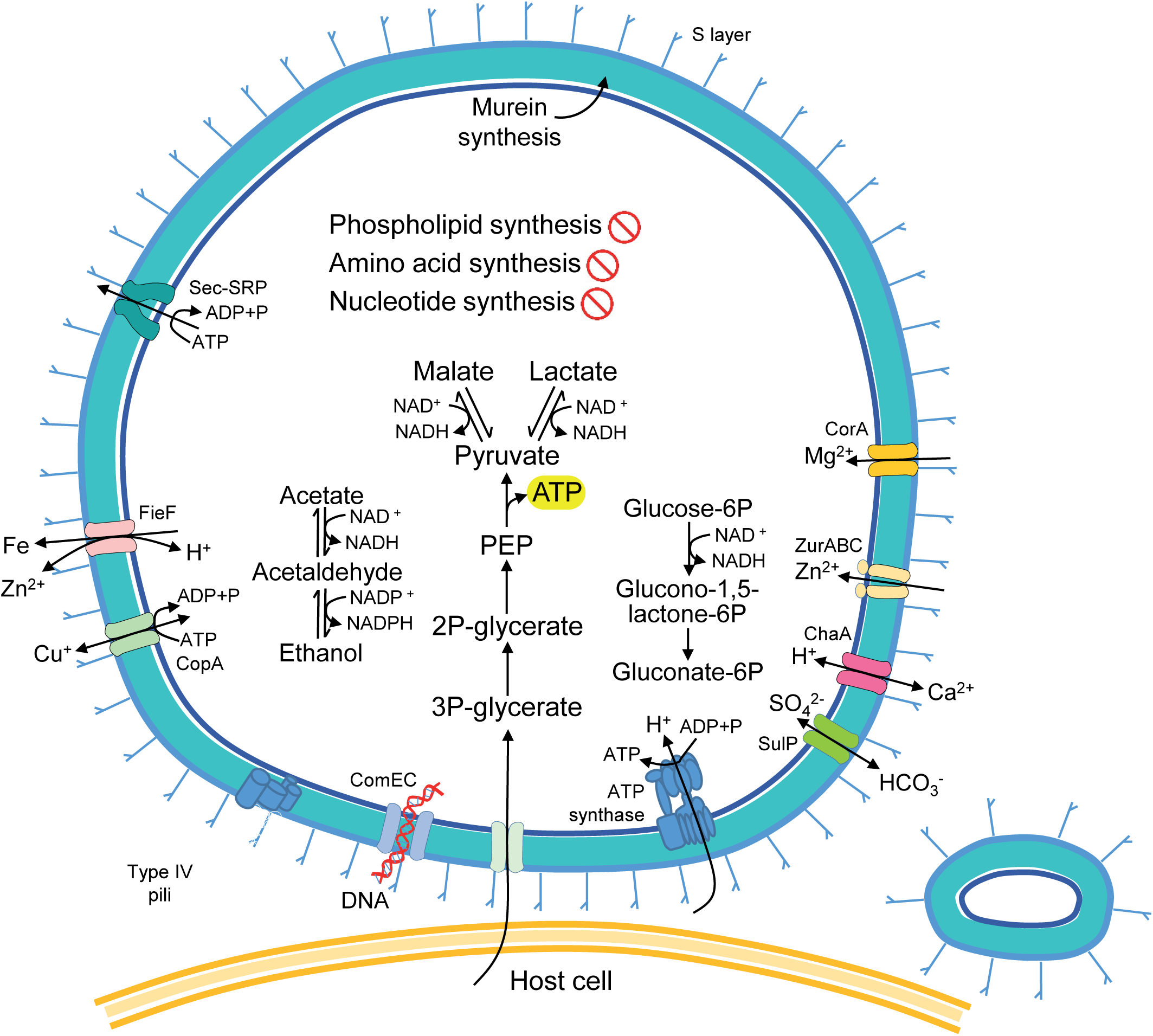
Reconstruction of metabolic capacities and structural features of *Strigamonas methylophilicida*. The diagram shows the host cell surface (bottom) with a *Strigamonas* cell attached to its surface and an extracellular vesicle on the right.

The *S. methylophilicida* MAGs encoded several additional carbon metabolism enzymes that are absent in the other characterized Absconditicoccaceae, including malate dehydrogenase, alcohol dehydrogenases, and aldehyde dehydrogenases. Thus, *S. methylophilicida* has the potential to produce or use malate, lactate, ethanol, and acetate from/to pyruvate (Fig. S10). These compounds have been hypothesized to be exported by Patescibacteria and used by other organisms, including their hosts (8). This intriguing possibility might suggest that some Patescibacteria can provide, at least under specific conditions, some benefit to their hosts. So far, patescibacterial symbiotic interactions seem to be parasitic, but whether they can occasionally and/or transiently become mutualistic (as is the case of some DPANN archaea (35)) remains to be shown (36). Additionally, *S. methylophilicida* encoded three enzymes of the pentose phosphate pathway: glucose-6-phosphate dehydrogenase and glucose-6-phosphate dehydrogenase, which both transform beta-D-glucose-6P into D-glucono-1,5-lactone-6P, and 6-phosphogluconolactonase, which further converts the latter product into D-gluconate-6P (Fig. 4 and Fig. S11). These reactions can generate NADPH, supplying the cell with reducing power.

The patescibacterial MAGs also encoded a complete F-type ATP synthase which, based on the sequence characteristics of its subunit c, probably translocates H^+^, using a proton motive force (PMF) to synthesize ATP, as suggested for *Vampirococcus* (14). However, similarly to many Patescibacteria (8), no classical electron transport chain was present. Alternative proteins to generate PMF proposed in the other two Absconditicoccaceae representatives (14, 15) were missing in *S. methylophilicida.* In contrast with the relatively large physical distance between cells of *Vampirococcus* and its host (∼100 nm), the cells of *S. methylophilicida* were more tightly attached to their host (∼20 nm), opening the possibility of direct salvage of protons from the host, as previously hypothesized for other Patescibacteria (8).

The *S. methylophilicida* MAGs encoded a relatively complex repertoire of cell surface components, a common trait in the family Absconditicoccaceae (14, 15). They contained genes for the type IV pilus (*pilBCDT*), probably involved in the attachment to the host, the natural competence system (*comEC*), probably related to DNA uptake from the host, and the secretion system Sec-SRP (*secDEFGYA* and *yidC*). As in *A. praedator* (15), we found diverse genes (10 in Smet_RSA and 8 in Smet_MG) coding for proteins with S-layer homology (SLH) domains. The S-layer is likely involved in the physical contact with the host cells. The complete peptidoglycan biosynthesis pathway was also present, as well as many ion channels and electrochemical potential-driven transporters. These transporters included Ca^2+^/H^+^ antiporters, Mg^2+^ transporters, Ca^2+^-activated Cl^-^ channels, SO_4_^2-^ permeases, and P-type Cu^+^ transporters. In addition, our MAGs encoded many ABC transporters (28 in RSA and 31 in MG). While many of these transporters could not be related to a particular function, others were inferred by sequence similarity to participate in cell division (*ftsX* and *ftsE*), DNA metabolism regulation (*upp*), and transport of metals such as Zn (*znuABC*). Smet_RSA and Smet_MG also encoded efflux systems for cobalt-zinc-cadmium and ferrous iron, and a *tetA* transporter of the Major Facilitator Superfamily (MFS), involved in the tetracycline efflux resistance mechanism.

### Cell defense mechanisms

The two *S. methylophilicida* MAGs encoded several DNA repair systems, including base excision, nucleotide excision, and mismatch repair systems (Fig. S12). Interestingly, the gene *mutT*, involved in the hydrolysis of 8-oxoguanine formed under oxidative stress (37), was present in eight copies, suggesting its particular importance in maintaining genome integrity. Other genes related to oxidative stress response were also detected, including those encoding glutathione peroxidase, glutathione *S*-transferase, thioredoxin reductase, thioredoxin-dependent peroxiredoxin, and superoxide dismutase Fe-Mn family members. Type I restriction-modification defense systems (*hsdRSM*) were present in four and two copies in Smet_RSA and Smet_MG, respectively. Both MAGs also encoded the type IV restriction enzyme Mrr but not the corresponding modification enzyme, whereas a cytosine-specific DNA methyltransferase was present in Smet_RSA. Both MAGs also encoded the virulence-related HigA-HigB and AbiEii-AbiEi toxin-antitoxin and abortive infection systems, which can be involved in the interaction of *S. methylophilicida* with its host and/or participate in antiviral defense through programmed cell death (38, 39). The toxin YafQ, an endoribonuclease that associates with the ribosome and blocks translation elongation through sequence-specific mRNA cleavage (40), was also detected in these MAGs.

We identified class I and class II CRISPR-Cas systems in *S. methylophilicida* (Fig. 5a) but, unexpectedly, neither of them included a complete adaptation module, which typically consists of *cas1*, *cas2*, and *cas4* genes. We only detected *cas4*, and only in Smet_RSA. The class I CRISPR-Cas system (41) was very partial, only represented by *cas6*, found in both *S. methylophilicida* MAGs, and *cas3*, found only in Smet_RSA. Neither other *cas* genes nor CRISPR arrays were found in the surrounding genomic context. Additionally, Smet_MG encoded *cas12a*, a signature of the class II type V CRISPR-Cas systems (42, 43). Similar CRISPR-Cas systems composed only of a CRISPR array and the *cas12a* gene have been previously reported (43). Cas12a is a multidomain enzyme that carries out the maturation of the pre-CRISPR RNA and the complete interference stage leading to the cleavage of the target DNA (43). The Smet_MG Cas12a was very divergent from those available in public databases. In particular, we only detected a fraction of the RuvC-like nuclease domain using the predicted 3D structure of the protein (Fig. S13). We identified two sequences highly similar to this divergent Cas12a encoded in two patescibacterial MAGs available in GenBank (JABCPD020000008.1, JAQVDN010000005.1; Fig. S14). However, in these patescibacterial MAGs, the CRISPR-Cas systems appear complete, including the adaptation module (Fig. 5a).

**FIG 5.**
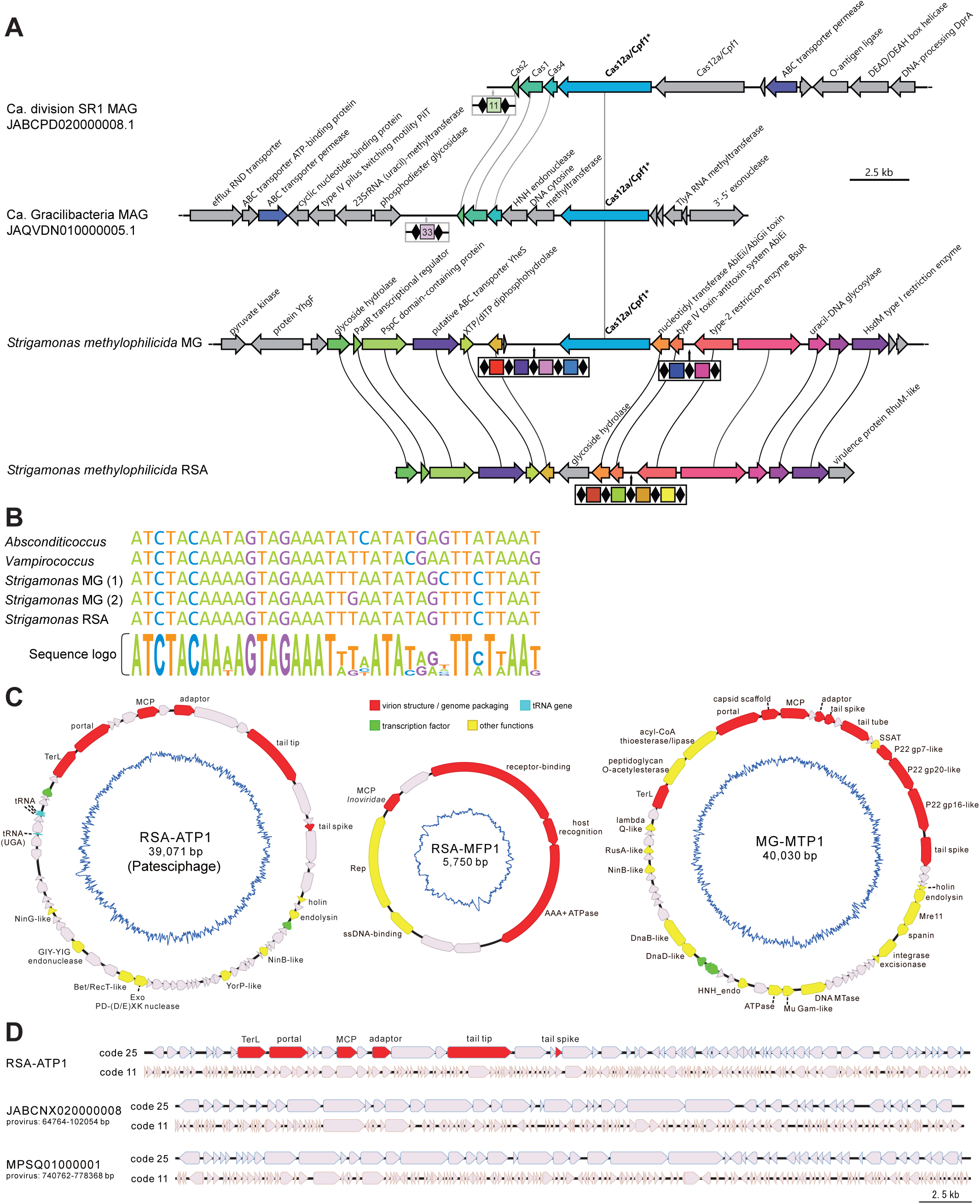
The CRISPR-Cas systems of *Strigamonas methylophilicida* and complete genomes of phages associated with Absconditicoccaceae and Methylophilaceae hosts. (A) CRISPR-Cas gene organization in the *S. methylophilicida* MAGs and two patescibacterial genomes with systems encoding similar Cas12a proteins. Spacers are represented as colored squares separated by direct repeats (black diamonds). ORFs shared by different taxa are colored. (B) Sequence alignment of the direct repeats of *S. methylophilicida* and other Absconditicoccaceae. A sequence logo under the alignment shows the conservation of each position. (C) Circular genome maps of the phages RSA-ATP1 (infecting *S. methylophilicida*) and RSA-MFP1 and MG-MTP1 (infecting the Methylophylaceae hosts). Genes responsible for virion morphogenesis are shown in red; tRNA genes in cyan; transcription factors in green; other functionally annotated genes are colored yellow; genes with unknown functions are shown in grey. The inner blue lines show the GC skews of the genomes. Phage names and genome lengths are indicated in the center of each map. Abbreviations: MCP, major capsid protein; Rep, rolling circle replication initiation endonuclease; SSAT, spermidine/spermine-N(1)-acetyltransferase; Mre11, phosphodiesterase/nuclease Mre11; DNA MTase, DNA methyltransferase; HNH_endo, HNH endonuclease; DnaD-like, DnaD-like replication initiation protein; DnaB-like, DnaB-like replicative helicase; NinB-like, NinB-like ssDNA-binding protein; RusA-like, RusA-like crossover junction endodeoxyribonuclease; TerL, terminase large subunit; NinG-like, NinG-like recombination protein. (D) Genome diagrams of RSA-ATP1 and two related prophages with genes predicted using code 25 (top) and 11 (bottom). RSA-ATP1 genes involved in virion morphogenesis are highlighted in red. The two prophages of Absconditabacteria are indicated with the GenBank accession numbers and the exact nucleotide coordinates.

Despite the differences mentioned above, the genomic region encoding the CRISPR-Cas systems presented conserved synteny in Smet_MG and Smet_RSA. Within this region, we identified CRISPR arrays with 4 spacers in Smet_RSA and 6 spacers in Smet_MG (Fig. 5a). These spacers showed no sequence similarity between the two MAGs, indicating that the two strains have experienced different phage infection histories in their respective ecosystems. The CRISPR array of Smet_MG was located in the middle of a 19 kb contig, which reduced the chance of missing adjacent *cas* genes due to MAG incompleteness. The absence of the adaptation module suggests that this system is incapable of adding new spacers, which is consistent with the small number of spacers in these CRISPR arrays. This contrasts with the genomes of *V. lugosii* and *A. praedator*, which not only contain several complete CRISPR-Cas systems but also large CRISPR arrays with 20-35 spacers. Interestingly, the direct repeat sequences of our MAGs were similar to those of *V. lugosii* and *A. praedator* (Fig. 5b), suggesting that these CRISPR-Cas systems derive from a common ancestral Absconditicoccaceae. The limited number of CRISPR spacers, the absence of the *cas1-cas2* adaptation module in both *S. methylophilicida* MAGs, and the loss of *cas12a* in Smet_RSA, collectively suggest that this ancestral Absconditicoccaceae CRISPR-Cas system is undergoing degradation in *S. methylophilicida*. To compensate for such loss, *S. methylophilicida* seems to have gathered in the nearby genomic context other defense systems, including restriction modification enzymes and an abortive infection (Abi) toxin-antitoxin system. This clustered organization in ‘defense islands’ is well known in bacteria (44).

### Identification of phages targeting *S. methylophilicida* and its host

To identify phages potentially associated with *Strigamonas* and its Methylophilaceae hosts, we analyzed the RSA and MG mini-metagenomes using geNomad (45). These mini-metagenomes yielded 45 and 39 putative viral contigs larger than 5 kb, respectively, with two (5,750 bp and 39,071 bp) and one (40,030 bp) being predicted as complete viral genomes assembled as circular DNA molecules (Fig. 5c). Analysis of the viral contigs using iPHoP, a computation pipeline that integrates multiple host prediction methods (46), did not yield reliable results. Thus, we focused on the three complete virus genomes for more detailed manual analysis. Notably, none of the phage genomes was targeted by CRISPR spacers identified in the *S. methylophilicida* MAGs, consistent with these CRISPR-Cas systems being inactive.

Considering that *S. methylophilicida* uses a modified genetic code (code 25), the open reading frames (ORFs) in the three virus genomes were called using both genetic codes 11 and 25. For one of the viral genomes (39,071 bp) from the RSA mini-metagenome (which we named RSA-ATP1, see below), signature viral proteins, such as portal and the large terminase subunit, could be predicted only when code 25 was used (Fig. 5d). In total, 60% (39/65) of ORFs in this phage genome contained internal UGA codons and, accordingly, were disrupted when translated using genetic code 11, strongly suggesting that the virus is associated with an Absconditicoccaceae host. Consistently, 11 RSA-ATP1 proteins yielded significant hits (>30% identity and E-value <1E-15) to Patescibacteria proteins in the NCBI nr database (see Materials and Methods and Table S3). Genomic neighborhood analysis of the corresponding hits allowed identification of two seemingly intact prophages, flanked by predicted attachment sites and integrated in the genomes of uncultured Absconditicoccaceae from the human oral microbiota (MPSQ01000001 and JABCNX020000008; Fig. S15a). Notably, similar to RSA-ATP1, many genes in the two prophages contained the internal UGA codons and thus the corresponding proteins could be translated only using the modified code 25 (Fig. 5d). Thus, we conclude that RSA-ATP1 infects an Absconditicoccaceae host, likely *S. methylophilicida*, given the enrichment of this species in the RSA mini-metagenomes.

BLASTp searches queried with the protein sequences of the two other phages identified in the RSA and MG mini-metagenomes also yielded best hits to bacterial genomes and analysis of the corresponding loci revealed the presence of related prophages (Fig. S15b-c). The prophage closely related to the 5,750 bp genome was identified in the chromosome of *Methylophilus* sp. Leaf414 (NZ_LMQQ01000004; Fig. S15b), a member of the family Methylophilaceae. By contrast, prophages distantly related to the 40,030 bp genome were present in the chromosomes of *Pseudomonas chlororaphis* and *Sphingomonas oryzagri* (Fig. S15c), two proteobacterial species from the phylum Pseudomonadota, to which Methylophilaceae also belongs. Thus, it is likely that the two phages present in the RSA and MG mini-metagenomes are associated with the Methylophilaceae hosts.

Based on the encoded signature proteins, namely, large terminase subunit, portal protein, and HK97-fold major capsid protein, both the phage likely associated with *Strigamonas* and the larger phage potentially infecting Methylophilaceae species could be assigned to the class *Caudoviricetes.* Accordingly, we denote these phages Ru Sainte Anne Absconditicoccaceae tailed phage 1 (RSA-ATP1) and Mare Gabard Methylophilaceae tailed phage 1 (MG-MTP1), respectively. Notably, MG-MTP1 encodes homologs of the T7 and P22 tail proteins (Fig. 5c and Table S4), suggesting that it has a podophage morphology with a short tail, whereas the high divergence of the RSA-ATP1 proteins makes more difficult to predict the tail phenotype of RSA-ATP1. The Methylophilaceae phage with the small circular genome encodes signature proteins conserved in filamentous phages of the order *Tubulavirales*; thus, we denote this phage as Ru Sainte Anne Methylophilaceae filamentous phage 1 (RSA-MFP1; Fig. 5c).

MG-MTP1 encodes an integrase and an excisionase and thus is likely to be a temperate virus, similar to the related prophages (Fig. S16). By contrast, neither RSA-ATP1 nor RSA-MFP1 encode detectable integrases, suggesting that neither of the two phages establish lysogeny. All filamentous phages are non-lytic and are released from the host cell through extrusion. By contrast, RSA-ATP1 encodes an endolysin and a potential holin, indicating that it is likely a lytic virus, despite the existence of related prophages in other patescibacterial genomes (Fig. S15a).

Only ∼30% of the RSA-ATP1 genes could be functionally annotated using sensitive profile-profile comparisons with HHsearch, as compared to 74% and 89% in MG-MTP1 and RSA-MFP1, respectively (Table S4). Besides the core virion morphogenesis proteins, this virus encodes a number of proteins involved in homologous recombination, including homologs of phage lambda proteins NinB, NinG, Exo and Bet (47), echoing the abundance of DNA repair genes (some of them exhibiting multiple paralogs) in its predicted host (see above). Although no potential auxiliary metabolic genes could be detected in the RSA-ATP1 genome, we found that this virus carries four tRNA genes. Remarkably, in addition to tRNA-Met(CAT), tRNA-Arg(TCT) and tRNA-Leu(TAA), RSA-ATP1 encodes the supressor tRNA-Gly^sup^(TCA), which reassigns the UGA stop codon to glycine. All four codons recognized by the phage-encoded tRNAs are used by both the host and the phage (Table S5) and thus the tRNAs may facilitate translation of the viral and cellular proteins. Notably, suppressor tRNAs have been previously described to be widespread in phages with large genomes (100-600 kb) and infecting bacteria with the standard genetic code (48). In these phages, suppressor tRNAs are thought to ensure timely expression of the late genes (48). However, to our knowledge, no patescibacterial phages with tRNA-Gly^sup^(TCA) genes were reported thus far. Notably, the two prophages detected in *Strigamonas* lack tRNA genes and hence likely depend on the host pool of tRNAs. BLASTtn search queried with the sequence of tRNA-Gly^sup^(TCA) against the NCBI non-redundant database and the IMG-VR database of virus genomes did not yield significant hits (>90% identity). Thus, the provenance of the suppressor tRNA gene in RSA-ATP1 remains unclear. Nevertheless, it is tempting to speculate that RSA-ATP1 and possibly certain other groups of patescibacterial phages have evolved from suppressor tRNA-encoding phages infecting other bacteria with the standard code.

To assess the novelty of the three phage genomes and their related prophages, we performed phylogenomic analyses using ViPTree (49), with the reference phage genomes available in the Virus-Host database (50) for comparison. Whereas RAS-MFP1 and the prophage of *Methylophilus* sp. Leaf414 were firmly placed within the *Inoviridae* family of filamentous phages, neither MG-MTP1 nor RSA-ATP1 were found to be closely related to any previously classified phages (Fig. S16). Thus, both tailed phages can be considered as founding representatives of two new families.

### Conclusions

The newly characterized species *Strigamonas methylophilicida* belongs to a novel genus in the patescibacterial family Absconditicoccaceae. This is the third genus in this family to undergo comprehensive characterization. The interaction between *S. methylophilicida* and Methylophilaceae hosts broadens the spectrum of known hosts of Patescibacteria. Like other Absconditicoccaceae, *S. methylophilicida* exhibits an ectoparasitic lifestyle, evident in TEM observations illustrating its small cell size and close attachment to the host. In addition, its genome sequence also indicates a strong reliance on the host to obtain essential components, including fatty acids, amino acids, and nucleotides. It also displays a substantial presence of strain– and genus-specific genes, supporting the previously reported dynamic evolution of gene content in the Absconditicoccaceae (14). *Strigamonas* stands out due to its large genome size, ∼1.9 Mb, and the production of extracellular vesicles, structures previously unseen in Patescibacteria and that could potentially be involved in the interaction with its host.

We unveiled novel phages that appear to target *S. methylophilicida* and its Methylophilaceae host. Two of the three phages for which complete genome sequences could be obtained represent new virus families within the class *Caudoviricetes*, whereas the third one represents a new species and potentially a new genus within the family *Inoviridae*. Similar to its predicted *S. methylophilicida* host, RSA-ATP1 uses a modified genetic code, suggesting adaptation to the host translation machinery. Notably, the phage encodes a tRNA which reassigns the stop codon UGA to glycine, a distinguishing feature of Patescibacteria. We also found that related prophages are integrated in the genomes of other Patescibacteria species, namely, the uncultured Absconditicoccaceae from the oral human microbiota. Thus, RSA-ATP1-related phages might represent a Patescibacteria-specific virus group, which apparently includes both temperate and lytic members.

### Taxonomy

Based on the genetic distance with other patescibacterial genomes, we propose that our new MAGs define two strains of a novel genus and species within the Absconditicoccaceae, which we have described under the SeqCode (51).

### Description of *Strigamonas* gen. nov

*Strigamonas* (Stri.ga.mo’nas. N.L. fem. n. *striga*, a female evil spirit or vampire; L. fem. n. *monas*, a unit, monad; N.L. fem. n. *Strigamonas*, a female vampire monad).

Epibiotic bacteria that parasitize on methylotrophic Gammaproteobacteria of the family Methylophilaceae. Non-flagellated cells attached to the surface of the host. Gram-positive cell wall structure.

The type species is *Strigamonas methylophilicida*.

### Description of Strigamonas methylophilicida sp. nov

*Strigamonas methylophilicida* (me.thy.lo.phi.li.ci’da. N.L. neut. n. *methylum*, the methyl group; from French masc. n. *méthyle*; from French masc. n. *méthylène*; from Gr. neut. n. *methy*, wine; from Gr. fem. n. *hylê*, wood; N.L. neut. n. *methyl*, pertaining to the methyl radical; N.L. masc. adj. suff. *-philus*, friend, loving; from Gr. masc. adj. *philos*, on; L. masc. n. suff. *-cida*, killer; from L. v. *caedo*, to cut, kill; N.L. masc. n. *methylophilicida*, a killer of the methyl radical lover).

Displays the following properties in addition to those given by the genus description. The cells are rounded and slightly flattened, approximately 300□nm in diameter, and form piles of up to 4 cells attached to the surface of their host. Found in freshwater environments.

It has a 1.9 Mb genome, with a G+C content of 27-28%. It lacks most of the biosynthetic pathways, most likely growing as an epibiotic parasite. It is known from environmental sequencing and microscopy observation only.

Smet_MG is the designated type MAG.

## MATERIALS AND METHODS

### Sampling and enrichment of patescibacteria-host consortia

Water samples were collected from two shallow freshwater ecosystems in the Regional Parc de la Haute Vallée de Chevreuse (south of Paris), including a small stream, Ru Sainte Anne (RSA), and a pond, Mare Gabard (MG) (23). We incubated water samples in 2 L glass bottles supplemented with 100 mM methanol at room temperature (RT). We regularly assessed the composition of the bacterial community using phase contrast light microscopy (see below). After approximately one month of incubation, we observed small cells attached to bigger, motile bacterial cells. We further enriched these consortia by inoculating 1 ml of the first enrichment in 5 ml of modified nitrate mineral salts (NMS) medium, known to promote the growth of methanotrophic and methylotrophic species (52). After one week of incubation at RT, we concentrated these consortia using fluorescence-activated cell sorting with a FACS Aria III flow cytometer (BD Biosciences, San Jose, CA, USA) equipped with one gas-state 633 nm laser and three solid-state 488 nm, 405 nm, and 375 nm lasers. Side and Forward Scatter Areas (SSC-A and FSC-A) were used to discriminate the epibiont-host consortia from other microbial cells. The 488 nm laser was used for the analysis of both FSC (488/10, 1.0 ND filter) and SSC (488/10) parameters applying voltages of 100V and 300V, respectively. The trigger was conjointly set to the FSC and SSC with a threshold of 500 and the fluidic system was run at 20 psi (3.102 bar) with a 100 µm nozzle. Samples were sorted at a speed of 3000–3500 events s^−1^ using sterile 1X PBS buffer as sheath fluid. We collected eight samples in 1.5 ml individual tubes (200 sorted events per tube) using the accurate “single-cell purity” sorting mode. The purity of the sorted events was visually inspected by phase contrast light microscopy (see below).

### Microscopy

Light microscopy observations of living cells were done with a Zeiss Axioplan 2 microscope equipped with a 100X oil-immersion phase contrast objective and a Nikon Coolpix B500 color camera for recording. For transmission electron microscopy (TEM), cells were fixed with 2.5% glutaraldehyde in 0.2M cacodylate buffer for 2 hours on ice. Then, cells were washed with 0.2M cacodylate, post-fixed in 1% osmium for 1 h at RT, and subsequently dehydrated with successive baths of increasing ethanol concentrations (30%, 50%, 70%, 80%, 90%, 95%, 2 x 100%, for 10 min each) and, finally, in a 1:1 100% ethanol:acetone mix for 10 min and twice in pure acetone for 10 min. The dehydrated cells were then embedded in a 1:1 acetone:epoxy resin mix overnight. Then, they were incubated in pure epoxy resin twice for 3 hours. Finally, the dehydrated cells were incubated overnight in pure epoxy resin at 70°C. Polymerized resin blocks were cut using a Leica EM UC6 ultramicrotome and sections were observed with a JEOL1400 microscope with an acceleration voltage of 80 KeV at the Imagerie-Gif facility (https://www.i2bc.paris-saclay.fr/bioimaging/).

### Individual cell isolation and 16S rRNA gene amplification and sequencing

Individual consortia of host-epibiont cells were collected with 6□µm-diameter microcapillaries (Eppendorf) mounted on an Eppendorf PatchMan NP2 micromanipulator set on a Leica Dlll3000 B inverted microscope. Cells were rinsed twice with sterile 10□mM Tris pH 8.0 buffer and finally deposited in a volume of 0.5□µl of this buffer for further processing.

DNA was purified from individual epibiont-host consortia with the PicoPure DNA extraction kit (Applied Biosystems). 16□S rRNA genes were amplified by PCR with the Platinum Taq DNA polymerase (Invitrogen) and the primers U515F (5’-GTGCCAGCMGCCGCGGTAA-3’) and 926R (5′-CCGYCAATTYMTTTRAGTTT-3′). PCR reactions consisted of 30 cycles (15□s denaturation at 94□°C, 30 s annealing at 55□°C, 2 min extension at 72□°C) preceded by 2□min denaturation at 94□°C, and followed by 7□min extension at 72□°C. Clone libraries of 16S rRNA gene amplicons were constructed with the Topo TA cloning system (Invitrogen) following the manufacturer’s instructions. After plating, positive transformants were screened by PCR amplification using the T7 (5’-TAATACGACTCACTATAGGG-3’) and M13 reverse (5’-CAGGAAACAGCTATGAC-3’) flanking vector primers. 16S rRNA gene amplicons were Sanger-sequenced using also using the T7 and M13R flanking vector primers by Genewiz (Essex, UK).

### Whole genome and 16S rRNA gene amplification

The REPLI-g Single Cell kit (Qiagen) was used to lyse and amplify whole genomes from RSA and MG FACS-sorted cells by multiple displacement amplification (MDA) following the manufacturer’s instructions. The amplified DNA products were diluted to 1/50. 16S rRNA genes were amplified by PCR with the Platinum Taq DNA polymerase (Invitrogen) and the following program: (i) 2 min initial denaturation at 94 °C, (ii) 35 cycles of 15 s denaturation at 94°C, 30 s annealing at 55 °C, and 90 s extension at 72 °C, (iii) 5 min final extension at 72 °C. The primers used were U515F (5’-GTGCCAGCMGCCGCGGTAA-3’) and 926R (5′-CCGYCAATTYMTTTRAGTTT-3′). 16S rRNA gene clone libraries were produced with the Topo TA cloning kit following the manufacturer’s instructions (Invitrogen). Individual clone inserts were amplified with vector primers T7 (5’-TAATACGACTCACTATAGGG-3’) and M13 reverse (5’-CAGGAAACAGCTATGAC-3’) and Sanger-sequenced by Genewiz (Essex, UK).

### Metagenome sequencing, MAG recovery, and functional annotation

We selected the five WGA (Whole Genome Amplification) products from the sorted RSA (three products) and MG (two products) cells that yielded 16S rRNA gene amplicons affiliating to Patescibacteria for direct Illumina HiSeq sequencing (2×125 bp paired-end reads; Eurofins Genomics, Ebersberg, Germany). Quality check and trimming of the reads were done with FastQC and Trimmomatic (MAXINFO:40:0.8, MINLEN:30 LEADING:20, TRAILING:20) (53, 54), respectively. Clean reads were assembled with SPAdes v3.15.3 (55) in MetaSPAdes mode using k-mer lengths of 21,25,31,35,41,45,51,55. To recover MAGs, we applied the anvi’o v6 metagenomic workflow (51) testing three different co-assembly strategies that yielded two high quality *S. methylophilicida* MAGS, Smet_RSA and Smet_MG. The Smet_RSA MAG was obtained by co-assembling the three RSA WGA clean reads; the Smet_MG MAG was derived from co-assembling the two MG WGA clean reads; and the 4 Methylophilaceae (host) MAGs were reconstructed by merging all the RSA and MG datasets. Within the anvi’o pipeline, we deliberately avoided using automated binning software to minimize the reliance on differential sequence coverage, which can be biased by the WGA. Contigs with a minimum length of 1000 bp were clustered hierarchically for manual binning via the anvi’o interactive interface (56). Completeness and redundancy of all MAGs were assessed with Anvi’o v6 (56). Taxonomic classification of the MAGs was performed with the GTDB-tk v2.3.0 pipeline (30, 57). We determined the phylogenetic novelty of Smet_RSA and Smet_MG by calculating their average amino acid identities against *V. lugosii* and *A. praedator* with the online AAI calculator (enve-omics.ce.gatech.edu/aai/) (58). Gene prediction for all MAGs was conducted with Prokka v1.14.5 (59) with annotations for the *Strigamonas* MAGs adjusted to accommodate codon usage 25. The resulting proteome was further annotated and visualized with the GhostKOALA web server (60). Protein structures were predicted with Alphafold (61). Structural comparisons to the Protein Data Bank (PDB) were done on the DALI server (62). We performed a gene orthology comparison among all the well-characterized Absconditabacterales MAGs (*Ca*. A. predator, *Ca*. V. lugosii, and our MAGs) with the anvi’o v7 pipeline (56, 63, 64). For these analyses, genes were compared in an all *vs*. all BLAST approach, clustered with MCL (65) (inflation parameter 2), annotated against the NCBI COG database (2014 release), and visualized using anvi’o v7.

### Phylogenetic analyses

16S rRNA gene sequences were aligned against the Silva 138.1 database with SINA (66). The multiple sequence alignment of the ∼100 closest sequences was trimmed with TrimAl v1.4.rev22-automated1 (67), and the corresponding maximum likelihood (ML) phylogenetic tree was reconstructed with IQ-TREE 1.6.11 (68) with the GTR+G model. Protein sequence alignments were produced using MAFFT v7.450-auto (69) and trimmed with trimAl v1.4.rev22-automated1 (67). Single-protein ML phylogenetic trees were constructed using IQ-TREE 1.6.11 (68) with the LG+C60+F+G model. For phylogenomic analyses, multiple sequence alignments were concatenated, and multi-protein ML phylogenetic trees were then reconstructed using IQ-TREE 1.6.11 with the LG+C60+F+G model. The gene markers used for the phylogenomic placement of the Patescibacteria were extracted using GTDB-tk, option identify (57). For all phylogenetic trees, branch supports were estimated using the ultrafast bootstrap method (1000 replicates) implemented in IQ-TREE 1.6.11 (68). Trees were visualized using iTOL (70).

### Identification and analysis of viral sequences

Putative viral sequences were extracted from metagenome assemblies using geNomad (github.com/apcamargo/genomad/) (45). Only complete genomes (i.e., sequences with direct terminal repeats or inverted terminal repeats) with virus scores over 0.70 were selected for subsequent analyses. The ORFs of the viral genomes were predicted using Pharokka (71), and functionally annotated using HHsearch (72) against the PFAM (Database of Protein Families), Protein Data Bank (PDB), Conserved Domain Database, and uniprot_sprot_vir70 databases. tRNA genes were predicted using ARAGORN (73) and tRNAscan-SE (74). Host prediction was attempted using iPHoP with default parameters (46). Sequences of terminase large subunits (TerLs), major capsid proteins (MCPs) and portal proteins of the identified complete viral genomes were used as BLASTp queries to search against the NCBI nr database. BLASTp hits with ≥30% identity and ≥50% coverage were retrieved and the corresponding sequences were downloaded for provirus search. Comparison and visualization of the viral genomes were carried out using Clinker (75). A genome-wide viral proteomic trees were generated using ViPTree web server (49).

### CRISPR spacer extraction and targeting

CRISPR-Cas loci in the bacterial MAGs were identified using CRISPRCasFinder (76). Then CRISPR spacers were extracted from all identified CRISPR-arrays and used to perform BLASTn searches against the MAGs and complete viral genomes. Sequence hits showing ≥95% identity and ≥95% coverage were considered as targets of the corresponding spacers.

## ACKNOWLEDGEMENTS

FB was supported by a PhD fellowship from the French Ministère de l’Enseignement Supérieur et de la Recherche. DM and PL-G were supported by grants from the European Research Council (ERC Advanced grants 787904 and 101141745, respectively) and by the ANR projects DArchFolds (ANR-22-CE02-0012-02) and ABiSYM (ANR-23-CE02-0016-01). LJ was supported by grants from CNRS-INSU-EC2CO and AGROPARISTECH. We thank the UNICELL platform (https://www.deemteam.fr/en/unicell) for help with FACS and single cell manipulation.

Study and experiment design: F.B., P.L.-G., P.D., D.M., L.J. Sampling and data collection: F.B., P.L.-G., A.G.-P., G.D., D.M., L.J. Optical and electron microscopy: F.B. Flow cytometry and molecular biology experiments: F.B., M.C., P.B., G.D. Patescibacteria, host, and phage genome sequence assembly, annotation, and interpretation: F.B., A.G.-P., M.K., Y.Z., P.L.-G., P.D., D.M., L.J. Manuscript writing: F.B., P.L.-G., M.K., D.M., L.J. Funding acquisition: L.J., P.L.-G., D.M. All authors read and approved the manuscript.

## DATA AVAILABILITY

The sequences generated in this study have been deposited in the Sequence Read Archive under Bioproject number PRJNA1086478.

## ADDITIONAL FILES

The following material is available online.

### Supplemental Material

Supplemental tables. Tables S1 to S15.

Supplemental material. Figures S1 to S16.

