## Supplementary figures for "Parasitic connections: a patescibacterial epibiont, its methylotrophic gammaproteobacterial host and their phages"

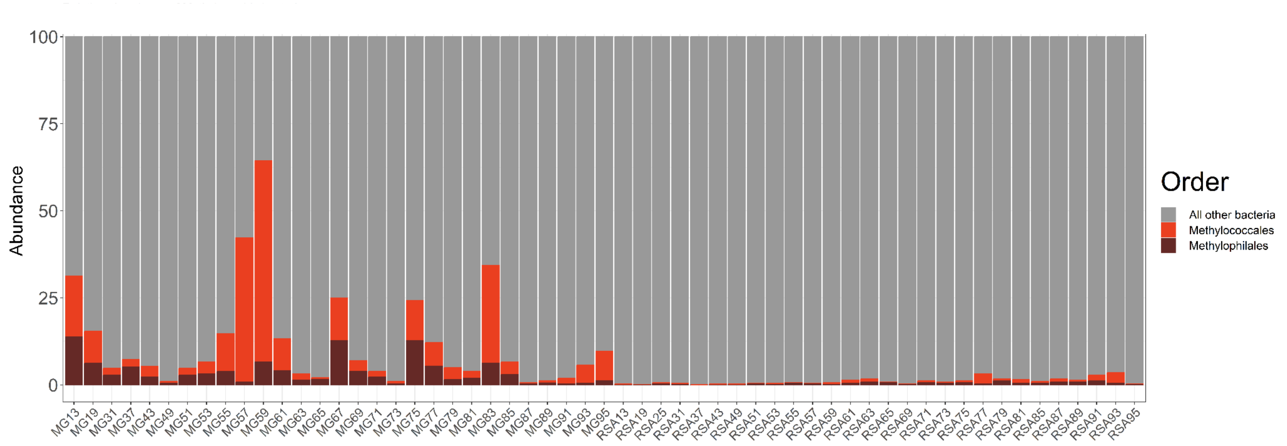


**FIG S1 Relative abundance of the orders Methylococcales and Methylophilales in the bacterial community of the Mare Gabard (MG) and Ru Sainte Anne (RSA) ecosystems**. These abundances are based on 16S rRNA gene metabarcoding analysis of samples collected every season between 2011 and 2019 (see David et al., 2021).

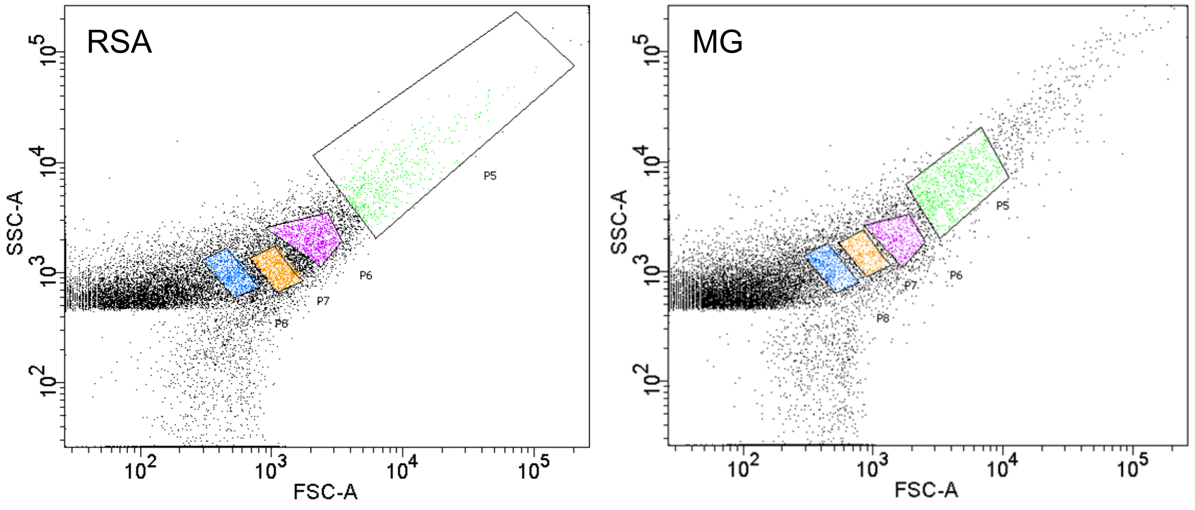


**FIG S2 Flow cytometry plots of the Side and Forward Scatter Areas (SSC-A and FSC-A, respectively) of enrichment samples from the Ru Sainte Anne (RSA) and Mare Gabard (MG) ecosystems**. Cell sorting gates are delimited by black lines and the detection events are colored differently. *Strigamonas methylophilicida* cells were found in sorted populations P6 (purple) and P7 (yellow).

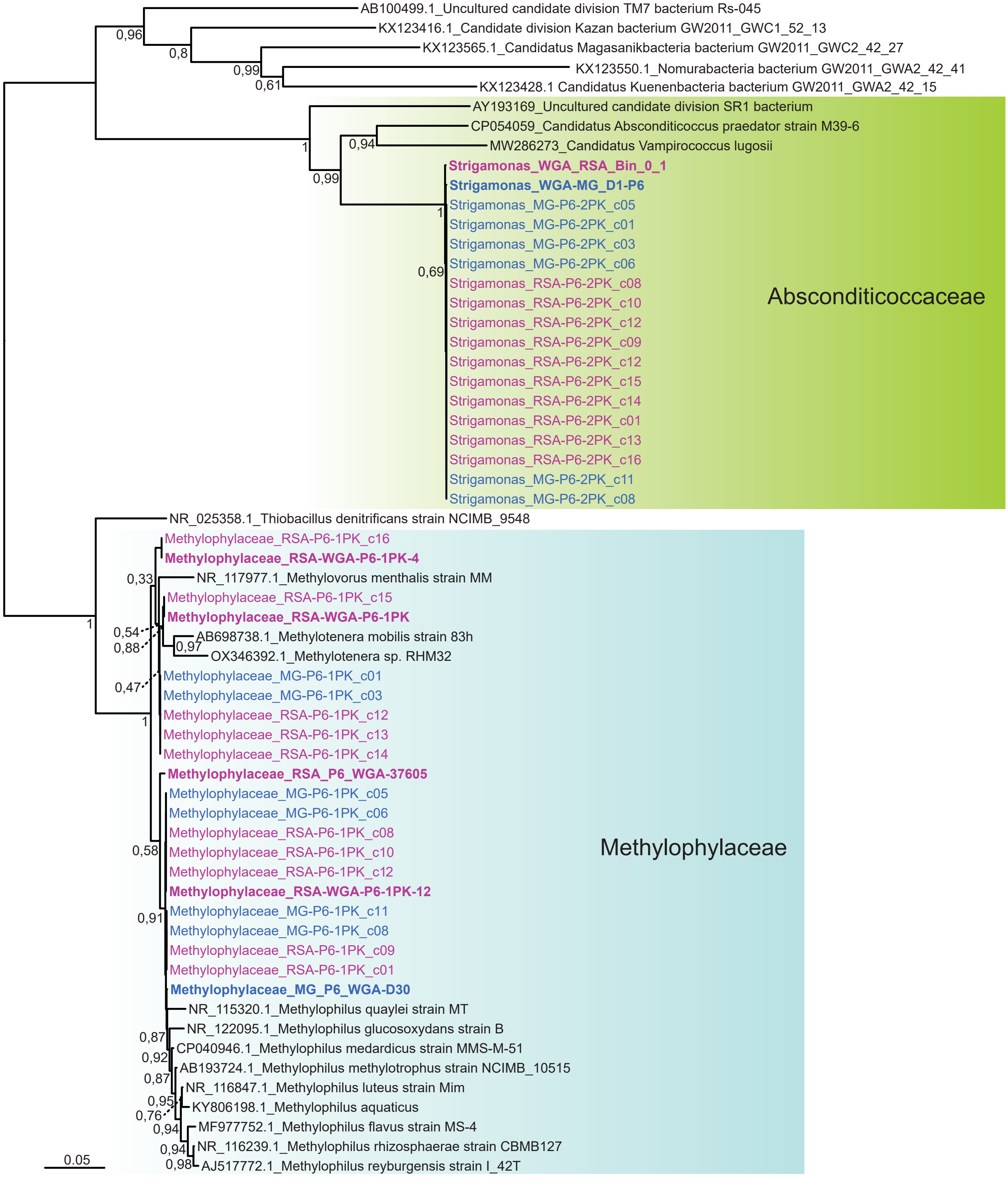


**FIG S3 Approximate maximum likelihood tree of 16S rRNA gene sequences obtained from single Abscondicoccaceae-host consortia**. Sequences of consortia obtained from the Mare Gabard (MG) ecosystem are in blue, and sequences of consortia obtained from the Ru Sainte Anne (RSA) ecosystem are in purple. Sequences in bold correspond to those obtained from massive sequencing after whole genome amplification (WGA) of DNA extracted from consortia sorted by flow cytometry. Numbers on branches indicate the statistical support.

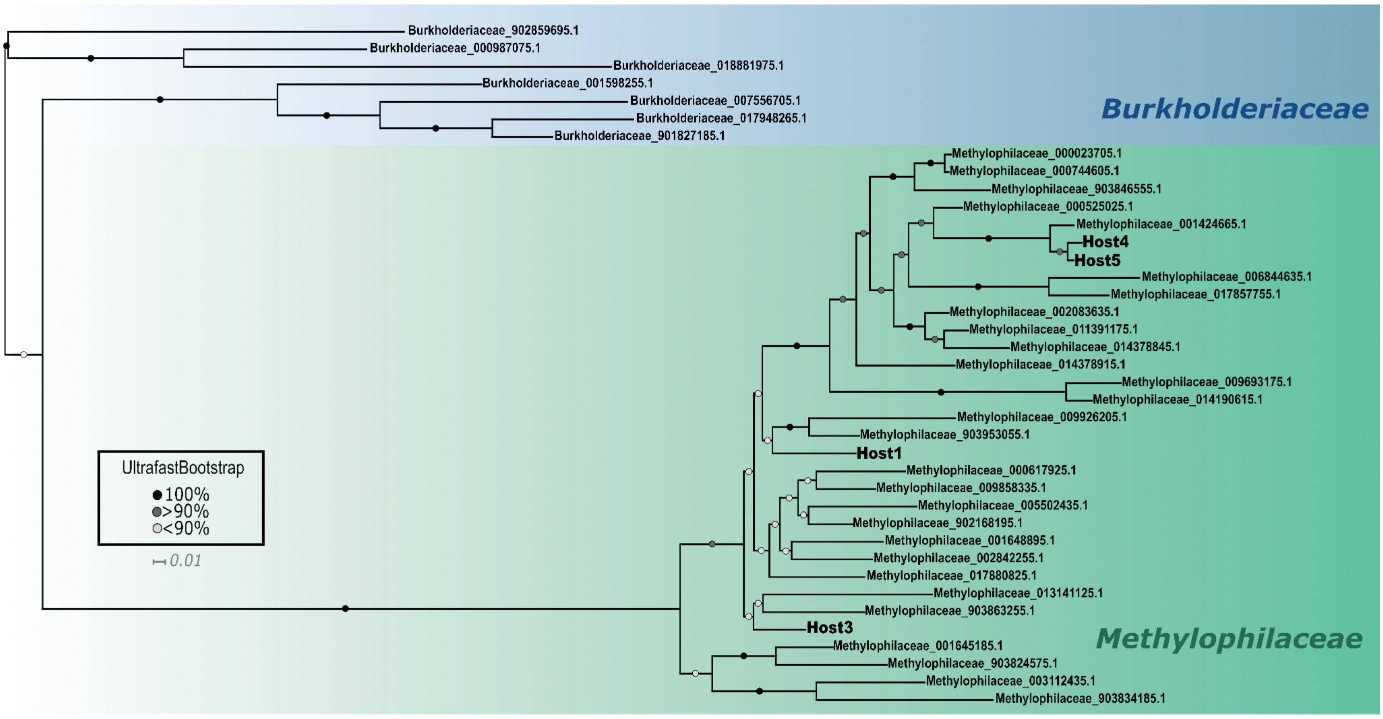


**FIG S4 Phylogenetic placement of the Methylophilaceae hosts of *Strigamonas*.** Maximum likelihood tree reconstructed using IQ-TREE v1.6.11 on a dataset of 14 concatenated ribosomal proteins (1900 aligned positions) using the LG+C60+F+G model. The four host MAGs (Host1, Host3, Host4, and Host5) branched within the family Methylophilaceae.

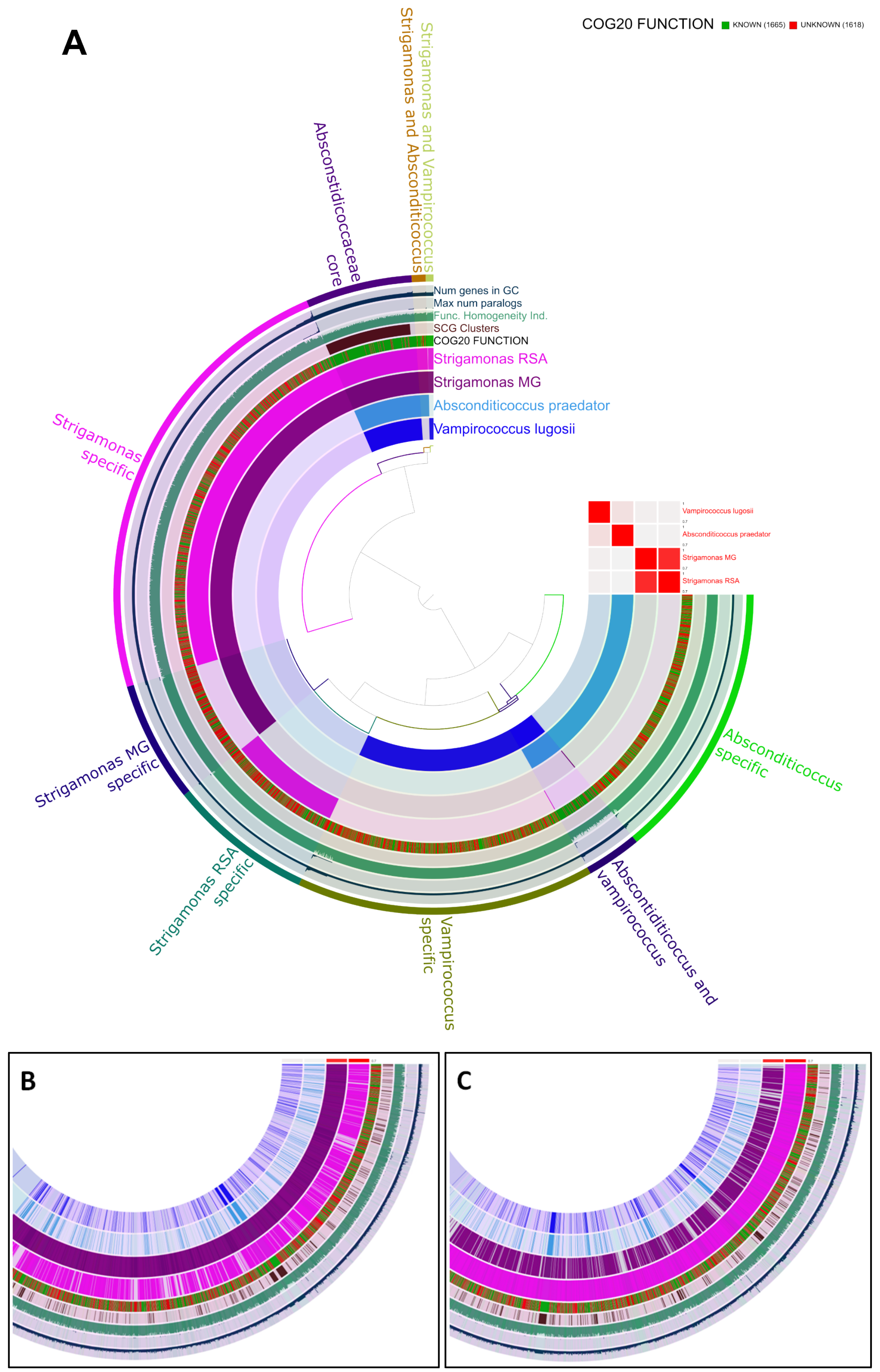


**FIG S5 Comparative genomics of the family Absconditicoccaceae. A.** Comparison of the inferred proteomes of our two *Strigamonas methylophilicida* MAGs (RSA in pink and MG in purple) with *Ca*. V. lugosii (dark blue) and *Ca*. A. praedator (light blue) (each of the four inner layers represents one MAG). Pangenomic representation from the Anvi'o pipeline showing the presence/absence of protein families, which are hierarchically clustered based on their presence-absence profiles in the different MAGs. The rest of the layers from inside to outside represent the COG functional annotation of each protein family (5th layer; known proteins in green; unknown in red); the families corresponding to Single Copy Genes (SCG) (6th layer); the functional homogeneity index of the protein family, which indicates the similarity in amino-acid composition of the proteins in each cluster (7th layer); the maximum number of paralogs per gene cluster (8th layer); The number of proteins per protein cluster (9$^{th}$ layer). The heatmap represents the ANI values between the four genomes. **B.** Zoom on a pangenome visualization where the gene families are re-organized based on the *S. methylophilicida* MG gene order. **C.** Zoom on a pangenome visualization where the gene families are re-organized based on the *S. methylophilicida* RSA gene order. Panels B and C show that many gene families uniquely present in either of the two *S. methylophilicida* MAGs are organized in clusters.

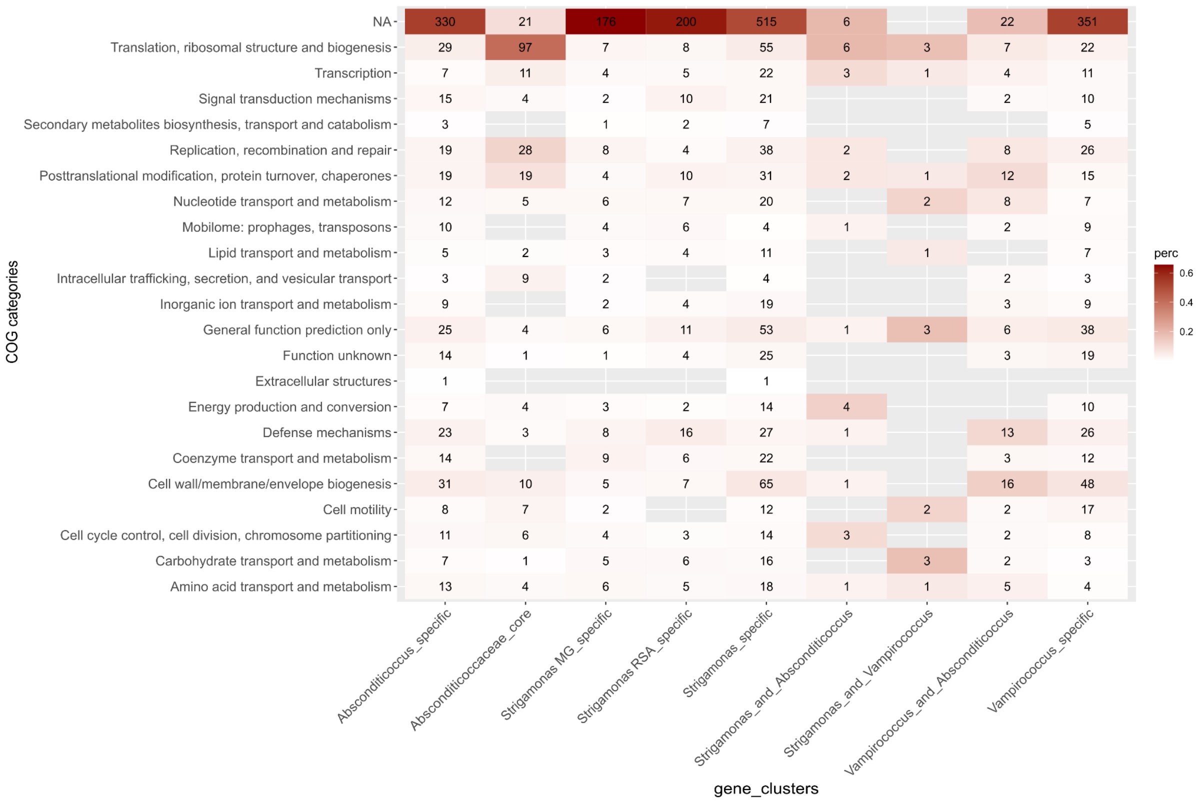


**FIG S6 Functional annotation of the family Absconditicoccaceae pangenome.** The heatmap shows the number of gene clusters with functional annotation based on COG categories shared between different members of the family Absconditicoccaceae. NA: not annotated.

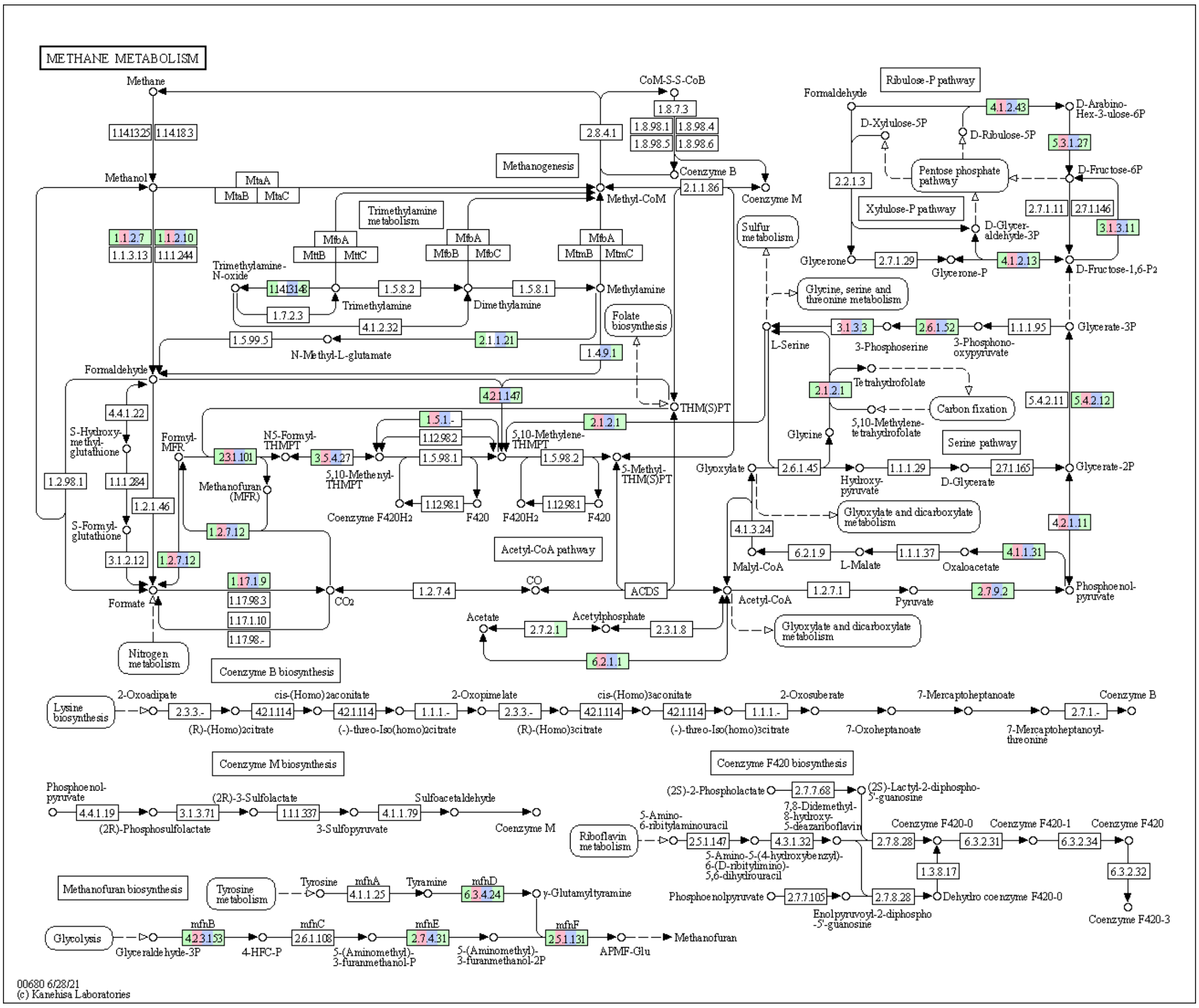


**FIG S7 KEGG map of the methane metabolism in the four Methylophilaceae host MAGs.** Boxes correspond to enzymes, when colored they indicate the presence of the corresponding gene in one of the four MAGs. Green, red, blue, and light green correspond, respectively, to host 4, host 5, host 3, and host 1.

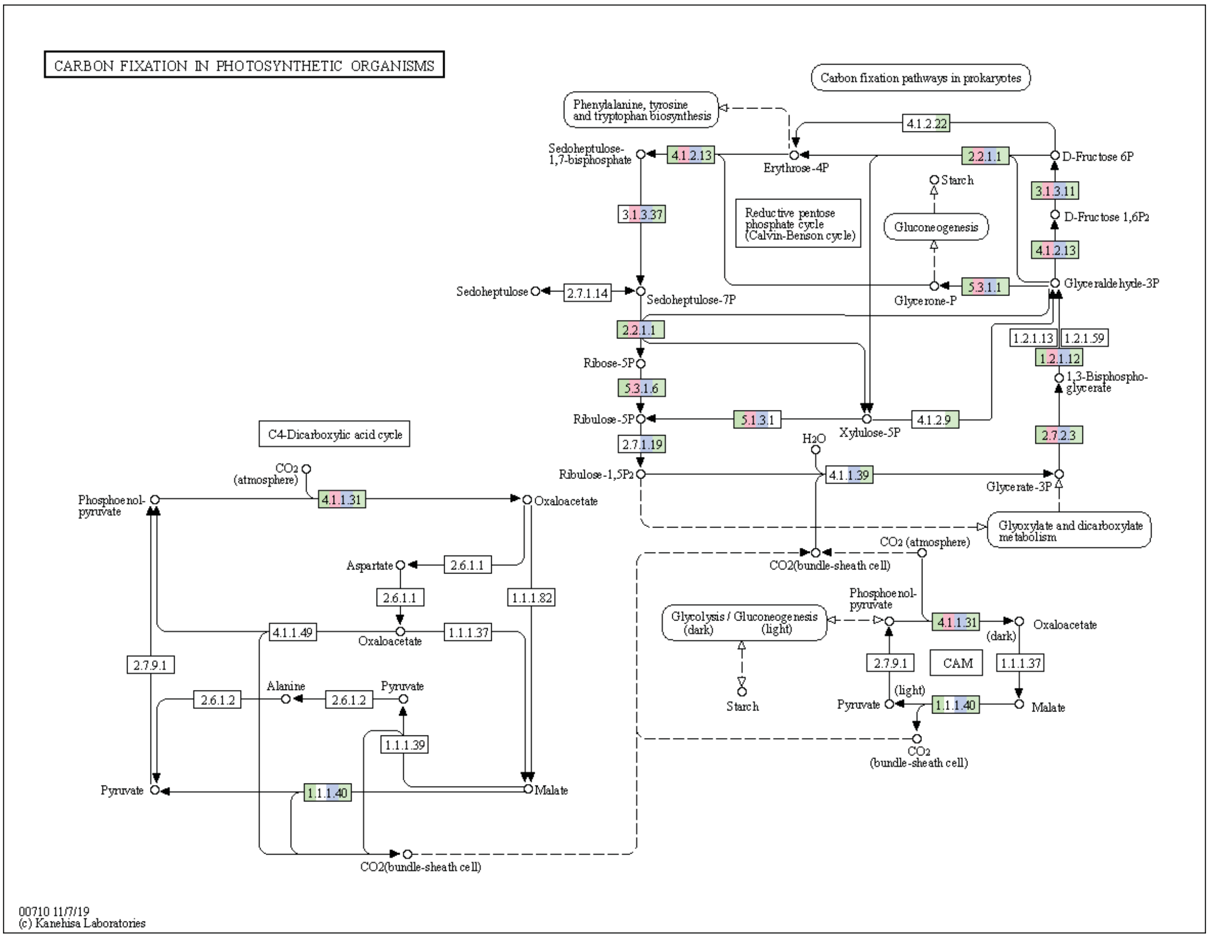


**FIG S8 KEGG map of the carbon fixation pathways in the four Methylophilaceae host MAGs**. Boxes correspond to enzymes, when colored, they indicate the presence of the corresponding gene in one of the MAGs. Green, red, blue, and light green correspond, respectively, to host 4, host 5, host 3, and host 1. RuBisCO (4.1.1.39) and PRK (2.7.1.19) were only found on MAGs of hosts 3 and 1.

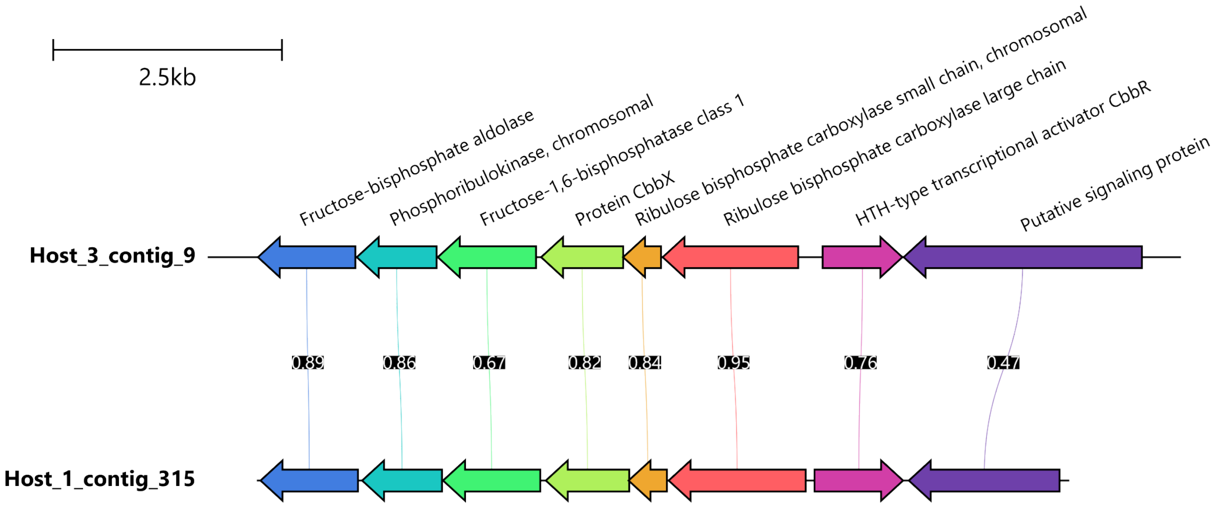


**FIG S9 Genomic context of the ribulose biphosphate carboxylase (RubisCO) genes in two Methylophilaceae host MAGs**. CDSs are represented with arrows, and their matching colors indicate sequence similarity between the MAGs (similarity values are shown in the black boxes between the similar ORFs).

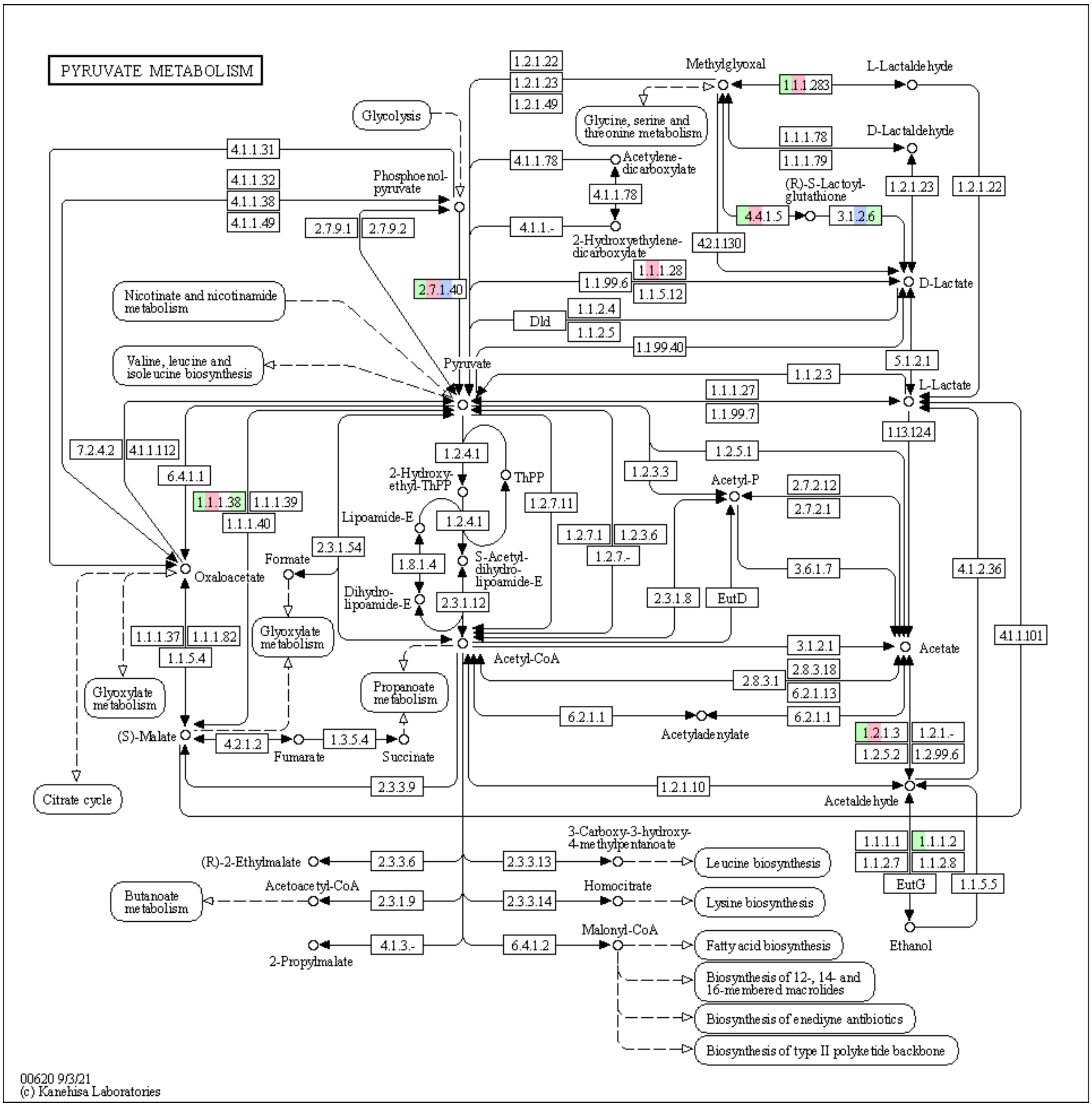


**FIG S10 KEGG map of the pyruvate metabolism in the Absconditicoccaceae**. Boxes correspond to enzymes, when colored, they indicate the presence of the corresponding gene in the four Absconditicoccaceae genomes: *S. methylophilicida* MG in green, *S. methylophilicida* RSA in red, *V. lugosii* in blue, and *A. praedator* in light green.

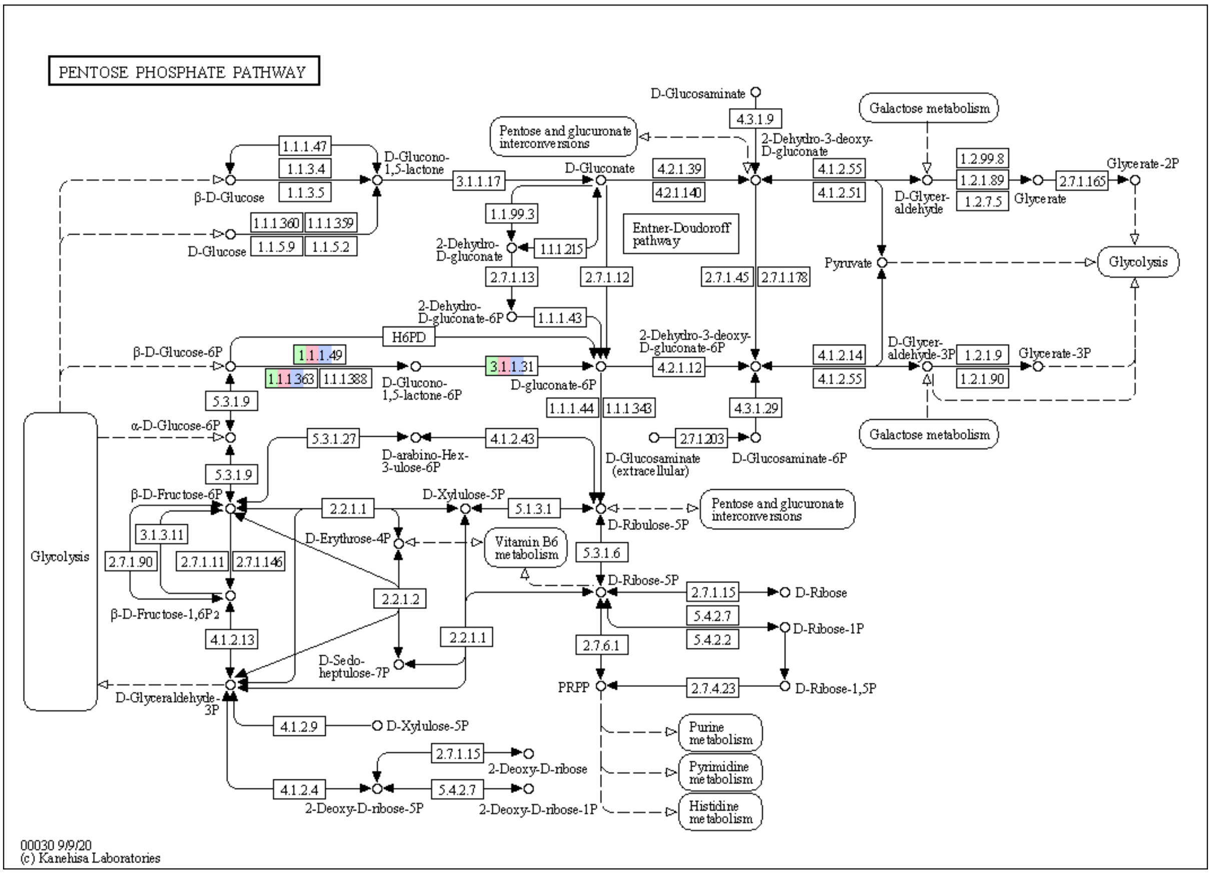


**FIG S11 KEGG map of the pentose phosphate pathway in the Absconditicoccaceae**. Boxes correspond to enzymes, when colored they indicate the presence of the corresponding gene in the four genomes: *S. methylophilicida* MG in green, *S. methylophilicida* RSA in red, *V. lugosii* in blue, and *A. praedator* in light green

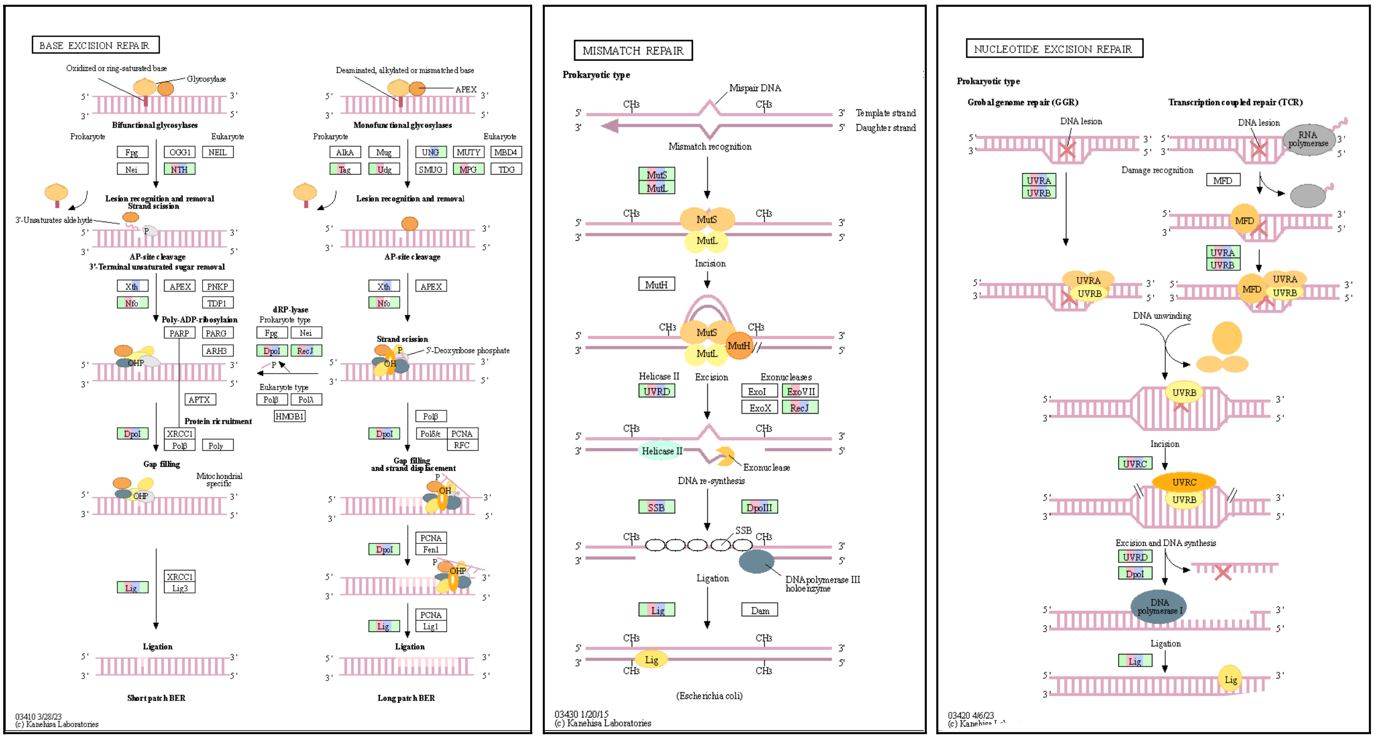


**FIG S12 KEGG map of the DNA repair systems in the Absconditicoccaceae**. Boxes correspond to enzymes, when colored, they indicate the presence of the corresponding gene: *S. methylophilicida* MG in green, *S. methylophilicida* RSA in red, *V. lugosii* in blue, and *A. praedator* in light green.

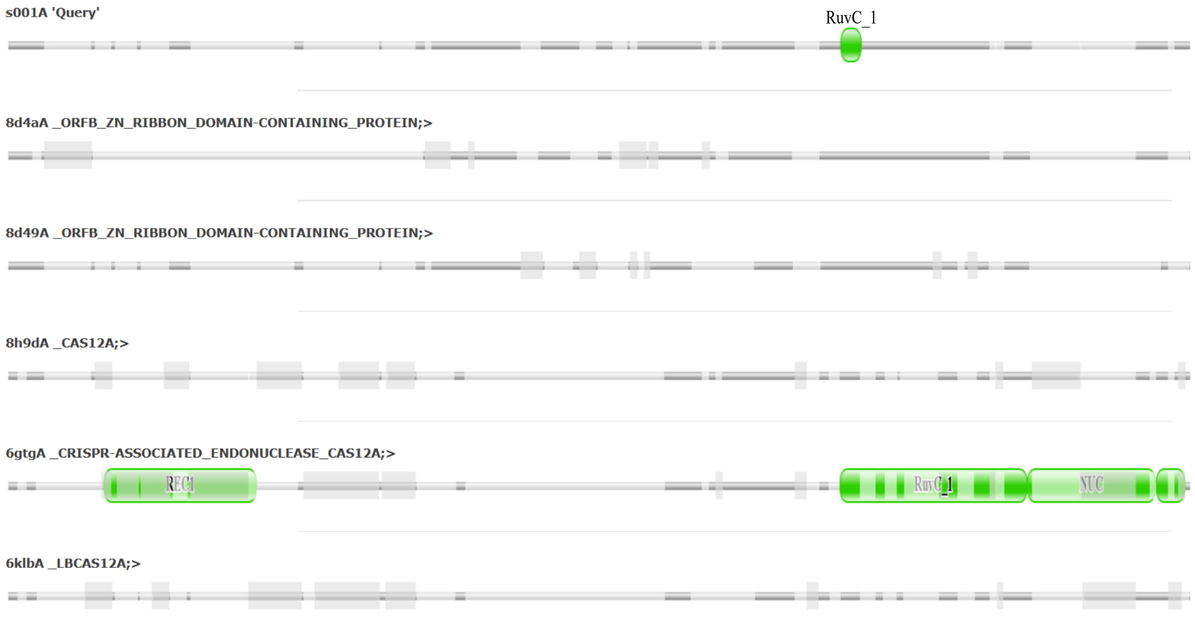


**FIG S13 Structural alignment of Cas12a-like proteins**. The alignment shows the *S. methylophilicida* MG Cas12a protein ('Query') aligned with the best structurally matching proteins (visualized with the DALI protein structure comparison server, http://ekhidna2.biocenter.helsinki.fi/dali/). Dark grey portions of the sequences indicate the regions structurally aligned with the *Strigamonas* sequence. Light grey portions indicate gaps added to the alignment, and thick grey portions indicate regions that are aligned among the other proteins but not with the *Strigamonas* one. Green blocks indicate sequences motifs annotated with Pfam v35.0 (released November 2021).

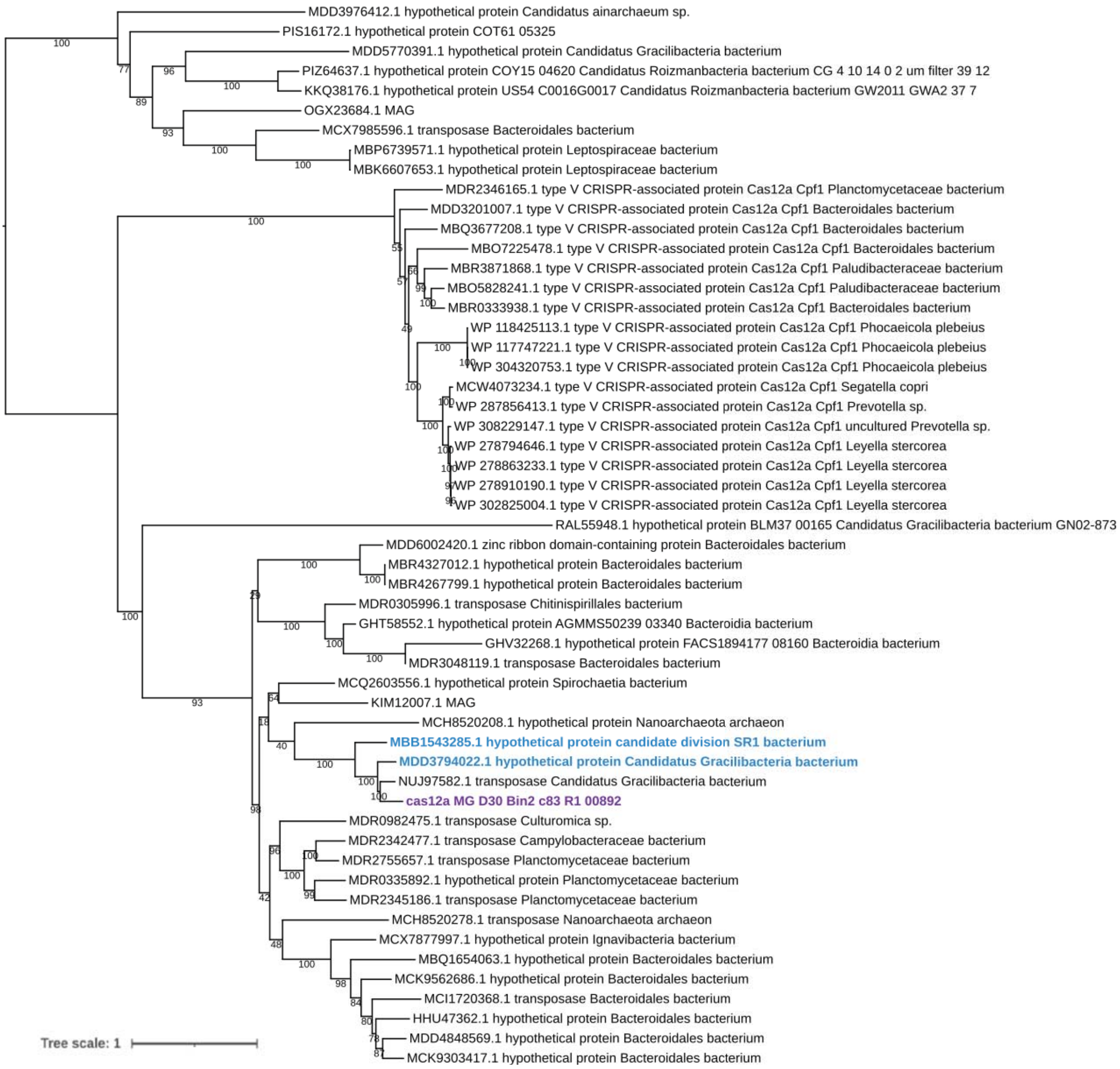


**FIG S14 Maximum likelihood phylogenetic tree of Cas12a**. The tree is based on an alignment of 1274 conserved amino acid positions. Support values on branches correspond to ultrafast bootstraps (1000 replicates). The sequence of *S. methylophilicida* MG is shown in purple. Two other Patescibacteria shown in blue contain a complete CRIPSR-Cas system in close proximity to the Cas12a coding gene (see Fig. 5).

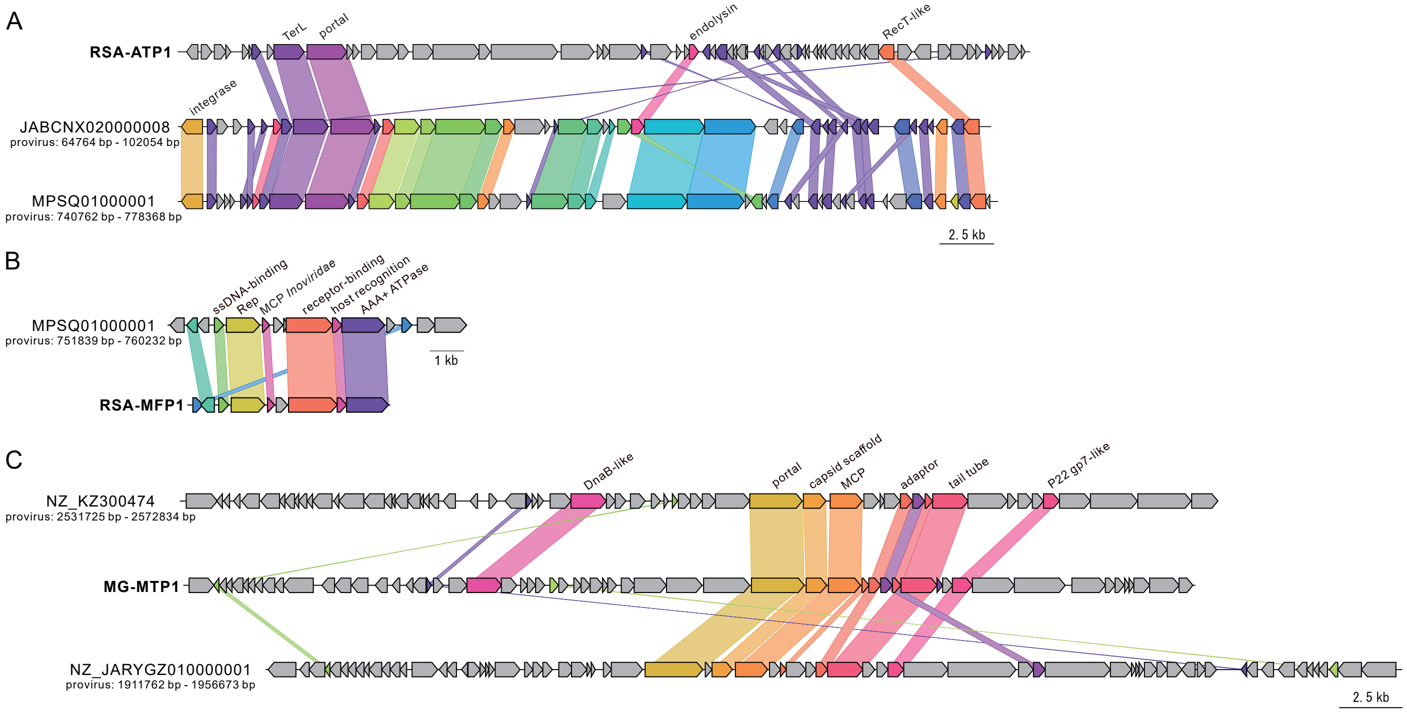


**FIG S15 Comparison of the Absconditicoccaceae and Methylophilaceae phages with related prophages.** (A) Gene content comparison between RSA-ATP1 and two related prophages of Patescibacteria found in the human oral microbiome. (B) Gene content comparison between RSA-MFP1 and a closely related prophage in the genome of *Methylophilus* sp. Leaf414. (C) Gene content comparison between MG-MTP1 and two closely related prophages of Pseudomonadota. All prophages are indicated with the corresponding genome accession numbers and exact nucleotide coordinates. Genes sharing over 20% identity are linked by shadings of the matching colors.

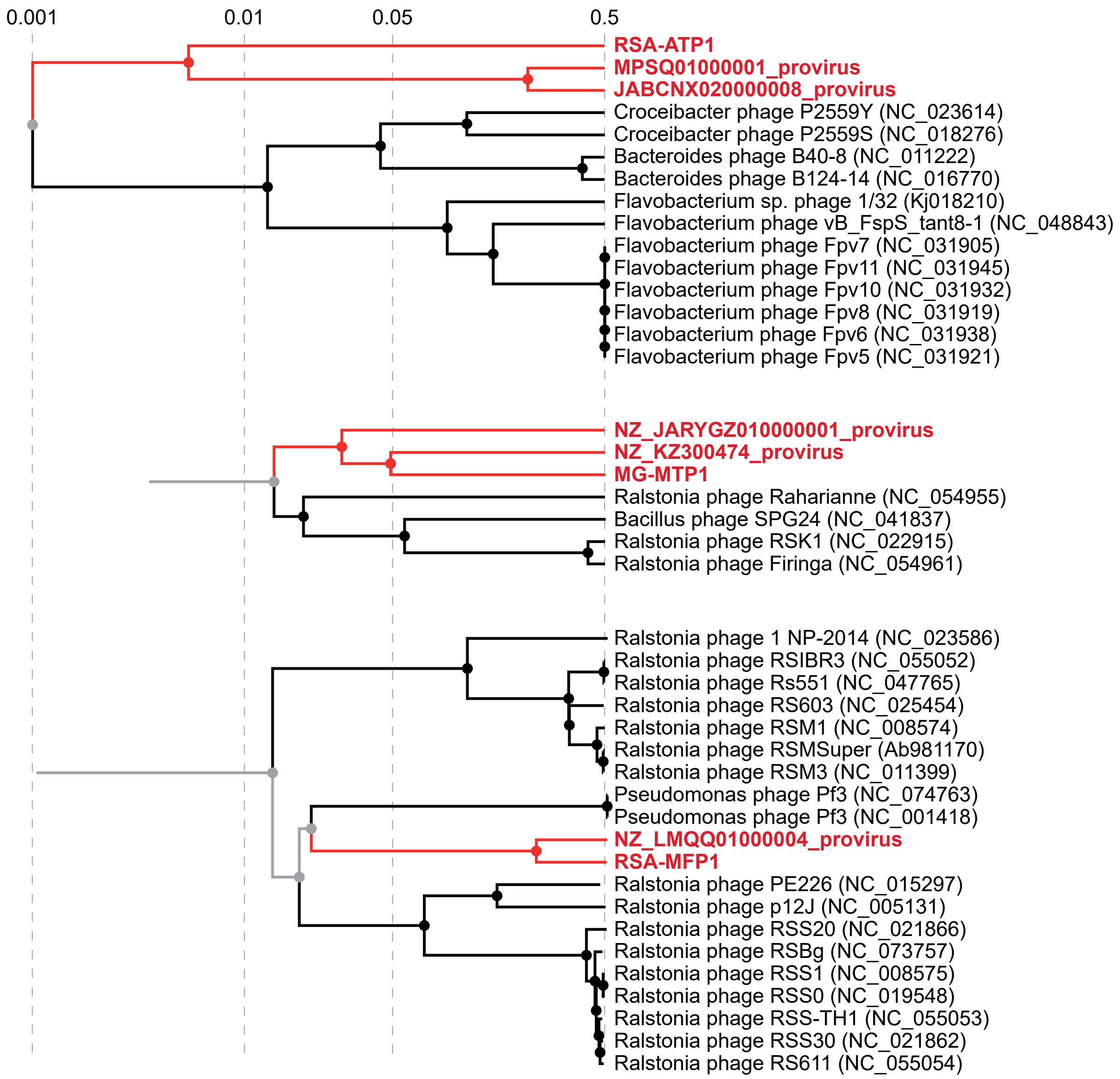


**FIG S16 Genome-wide proteomic trees of the phages RSA-ATP1, RSA-MFP1 and MG-MTP1**. Branches corresponding to the three phages and related prophages are highlighted in red. The trees are based on all-versus-all proteomic similarity matrices and mid-point rooted. Branch lengths are log-scaled. In the case of *Caudoviricetes*, the branch length of 0.05 is commonly taken as a family-level demarcation.
